# Identification and quantification of SARS-CoV-2 leader subgenomic mRNA gene junctions in nasopharyngeal samples shows phasic transcription in animal models of COVID-19 and dysregulation at later time points that can also be identified in humans

**DOI:** 10.1101/2021.03.03.433753

**Authors:** Xiaofeng Dong, Rebekah Penrice-Randal, Hannah Goldswain, Tessa Prince, Nadine Randle, Javier Salguero, Julia Tree, Ecaterina Vamos, Charlotte Nelson, James P. Stewart, ISARIC4C Investigators, COVID-19 Genomics UK (COG-UK) Consortium, Malcolm G. Semple, J. Kenneth Baillie, Peter J. M. Openshaw, Lance Turtle, David A. Matthews, Miles W. Carroll, Alistair C. Darby, Julian A. Hiscox

## Abstract

**Introduction:** SARS-CoV-2 has a complex strategy for the transcription of viral subgenomic mRNAs (sgmRNAs), which are targets for nucleic acid diagnostics. Each of these sgRNAs has a unique 5’ sequence, the leader-transcriptional regulatory sequence gene junction (leader-TRS-junction), that can be identified using sequencing.

**Results:** High resolution sequencing has been used to investigate the biology of SARS-CoV-2 and the host response in cell culture models and from clinical samples. LeTRS, a bioinformatics tool, was developed to identify leader-TRS-junctions and be used as a proxy to quantify sgmRNAs for understanding virus biology. This was tested on published datasets and clinical samples from patients and longitudinal samples from animal models with COVID-19.

**Discussion:** LeTRS identified known leader-TRS-junctions and identified novel species that were common across different species. The data indicated multi-phasic abundance of sgmRNAs in two different animal models, with spikes in sgmRNA abundance reflected in human samples, and therefore has implications for transmission models and nucleic acid-based diagnostics.

## Introduction

Various sequencing approaches are used to characterise SARS-CoV-2 RNA synthesis in cell culture^1, 2^, ex vivo models^3^ and clinical samples. This can include nasopharyngeal swabs from patients with COVID-19^4^ to post-mortem samples from patients who died of severe disease^5^. Bioinformatic interrogation of this data can provide critical information on the biology of the virus. SARS-CoV-2 genomes are message sense, and the 5’ two thirds of the genome is translated and proteolytically cleaved into a variety of functional subunits, many of which are involved in the synthesis of viral RNA^6^. The remaining one third of the genome is expressed through a nested set of subgenomic mRNAs (sgmRNAs). These have common 5’ and 3’ ends with the coronavirus genome, including a leader sequence. Many studies have shown that the sgmRNA located towards the 3’ end of the genome, which encodes the nucleoprotein, generally has a higher abundance than those located immediately after the 1a/b region and the genome itself^7, 8^. However, there is not necessarily a precise transcription gradient of the sgmRNAs. The 5’ leader sequence on the sgmRNAs is immediately abutted to a short sequence called a transcriptional regulatory sequence (TRS) that is involved in the control of sgmRNA synthesis^9, 10^. These TRSs are located along the genome and are proximal to the start codons of the open reading frames^11^. In the negative sense the TRSs are complementary to a short portion of the genomic leader sequence. The TRS is composed of a short core motif that is conserved and flanking sequences^9, 10, 12^. For SARS-CoV-2 the core motif is ACGAAC.

The prevailing thought is that synthesis of sgmRNAs involves a discontinuous step during negative strand synthesis^13, 14^. A natural consequence of this is recombination resulting in insertions and deletions in the viral genome and the formation of defective viral RNAs. Thus, the identification of the leader/sgmRNA complexes by sequencing provides information on the abundance of the sgmRNAs and evidence that transcription has occurred in the tissue being analysed. In terms of clinical samples, if infected cells are present, then leader/sgmRNA ‘fusion’ sequence can be identified, and inferences made about active viral RNA synthesis from the relative abundance of the sgmRNAs. In the absence of human challenge models, the kinetics of virus infection are unknown, and most studies will begin with detectable viral RNA on presentation of the patient with clinical symptoms. In general, models of infection of humans with SARS-CoV-2 assume an exponential increase in viral RNA synthesis followed by a decrease as antibody levels increase^15^.

In order to investigate the presence of SARS-CoV-2 sgmRNAs in clinical (and other) samples, a bioinformatics tool (LeTRS), was developed to analyse sequencing data from SARS-CoV-2 infections by identifying the unique leader-TRS gene junction site for each sgmRNA. The utility of this tool was demonstrated on cells infected in culture, nasopharyngeal samples from human infections and longitudinal analysis of nasopharyngeal samples from two non-human primate models for COVID-19. The results have implications for diagnostics and disease modelling.

## Results

A tool, LeTRS (named after the leader-TRS fusion site), was developed to detect and quantify defined leader gene junctions of SARS-CoV-2 (and other coronaviruses) from multiple types of sequencing data. This was used to investigate SARS-CoV-2 sgmRNA synthesis in humans and non-human primate animal models. LeTRS was developed using the Perl programming language, including a main program for the identification of sgmRNAs and a script for plotting graphs of the results. The tool accepts FastQ files derived from Illumina paired-end or Oxford Nanopore amplicon cDNA, Nanopore direct RNA sequencing, or BAM files produced by a splicing alignment method with a SARS-CoV-2 genome (Supplementary Figure 1). (Note that SARS-CoV-2 sgmRNAs are not formed by splicing, but this is the apparent observation from sequencing data as a result of the discontinuous nature of transcription). By default, LeTRS analyses SARS-CoV-2 sequence data by using 10 known leader-TRS junctions and an NCBI reference genome (NC_045512.2). However, given the potential heterogeneity in the leader-TRS region and potential novel sgmRNAs, the user can also provide customize leader-TRS junctions and SARS-CoV-2 variants as a reference. The tool was designed to investigate very large data sets that are produced during sequencing of multiple samples. As there is some heterogeneity in the leader-TRS sites, LeTRS was also designed to search the leader-TRS junction in a given interval, report 20 nucleotides at the 3’ end of the leader sequence, the TRS and translated first orf of the sgmRNA, and find the conserved ACGAAC sequences in the TRS.

### Combinations of read alignments with the leader-TRS junction that are considered for identifying leader-TRS junction sites

Various approaches have been used to sequence the SARS-CoV-2 genome and in most cases, this would also include any sgmRNAs as they are 3’ co-terminal and share common sequence extending from the 3’ end. Methods such as ARTIC^16^ and RSLA^4^ use primer sets to generate overlapping amplicons that span the entire genome, and also amplify sgmRNA. Included is a primer to the leader sequence, so that the unique 5’ end of these moieties are also sequenced. Primer sets of ARTIC and RSLA are generally pooled. Unbiased sequencing can also be used in methodologies to identify SARS-CoV-2 sequence. Data in the GISAID database have been generated by Oxford Nanopore (minority) or Illumina (majority) based approaches. These can give different types of sequencing reads derived from the sgmRNAs that can be mapped back on the reference SARS-CoV-2 genome by splicing alignment (Figure 1A). For example, there are a number of different types of reads that can be derived from mapping Illumina-based amplicon sequencing onto the reference viral genome (Figure 1B and 1C). During the PCR stage, the extension time allows the leader-TRS region on the sgmRNAs to be PCR-amplified by the forward primer and the reverse primer before and after leader-TRS junction in different primer sets, respectively. Both of these forward and reverse and primers would be detected at both ends of each paired read (Figure 1B pink lines) if the amplicon had a length shorter than the Illumina read length (usually 100-250 nts). If the amplicon was longer than the Illumina read length, primer sequence would be only found at one end of each paired read (Figure 1B green lines). The extension stage could also proceed with a single primer using cDNA from the sgmRNA as template. This type of PCR has a very low amplification efficiency, but theoretically could also generate the same Illumina paired end reads with just one primer sequence at one end (Figure 1C). These paired end reads could include the full length of the leader sequence but might not reach the 3’ end of the sgmRNA, because of the limitation of Illumina sequencing length and extension time (Figure 1C). Also, unless there are cryptic TRSs, all sgmRNAs would be expected to be larger than the Illumina sequencing length.

**Figure 1.**
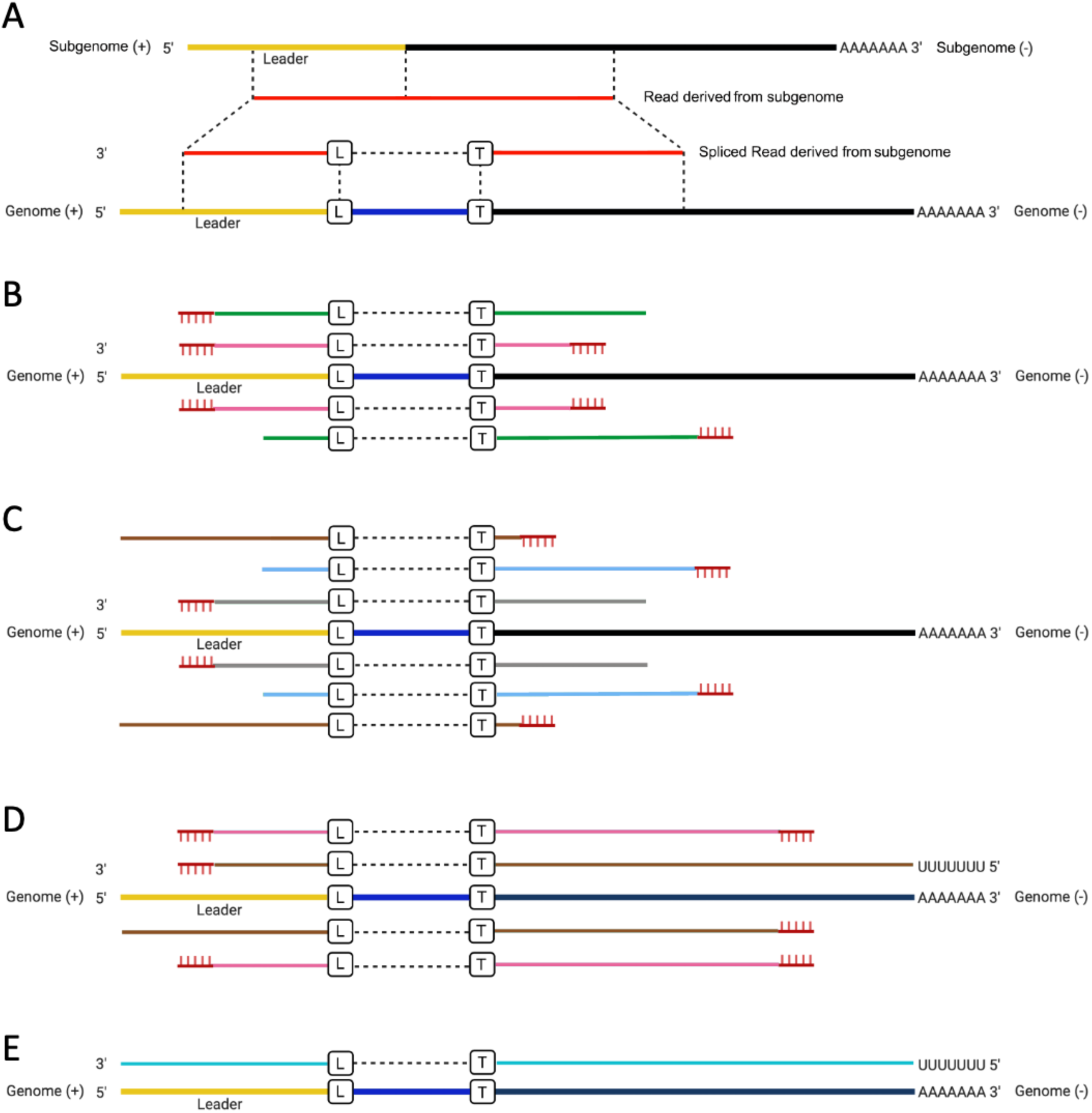
(A) Illustration of reads derived from sgmRNAs mapped onto the SARS-CoV-2 reference genome with a splicing method. Illustration of the possible type of reads mapped on the SARS-CoV-2 reference genome for the (B and C) paired end Illumina cDNA amplicon sequencing, where the lines with same colour implied paired reads, (D) Nanopore cDNA amplicon sequencing and (E) Nanopore direct RNA sequencing of SARS-CoV-2 genome. L and B in the boxes indicate the leader-TRS breaking sites on the leader side and TRS side, respectively.

In contrast, the different types of read alignment in the Nanopore based cDNA sequencing are simpler to assign. The longer reads that tend to be generated by Nanopore sequencing (depending on optimisation) enable the capture of full-length sequences of all amplicons. Provided the leader sequence is included as a forward primer most of the reads spanning the leader-TRS junction would contain the forward and reverse primer sequences at both ends (Figure 1D pink lines). If the extension time allowed, single primer PCR amplification could take the Nanopore cDNA sequencing reads to both the 3’ and 5’ ends of the sgmRNAs, and these types of reads would only have a primer sequence at one end (Figure 2D brown lines). In the Nanopore direct RNA sequencing approach, the full length sgmRNA could be sequenced and mapped entirely on the leader and TRS-orf regions (Figure 1E).

**Figure 2.**
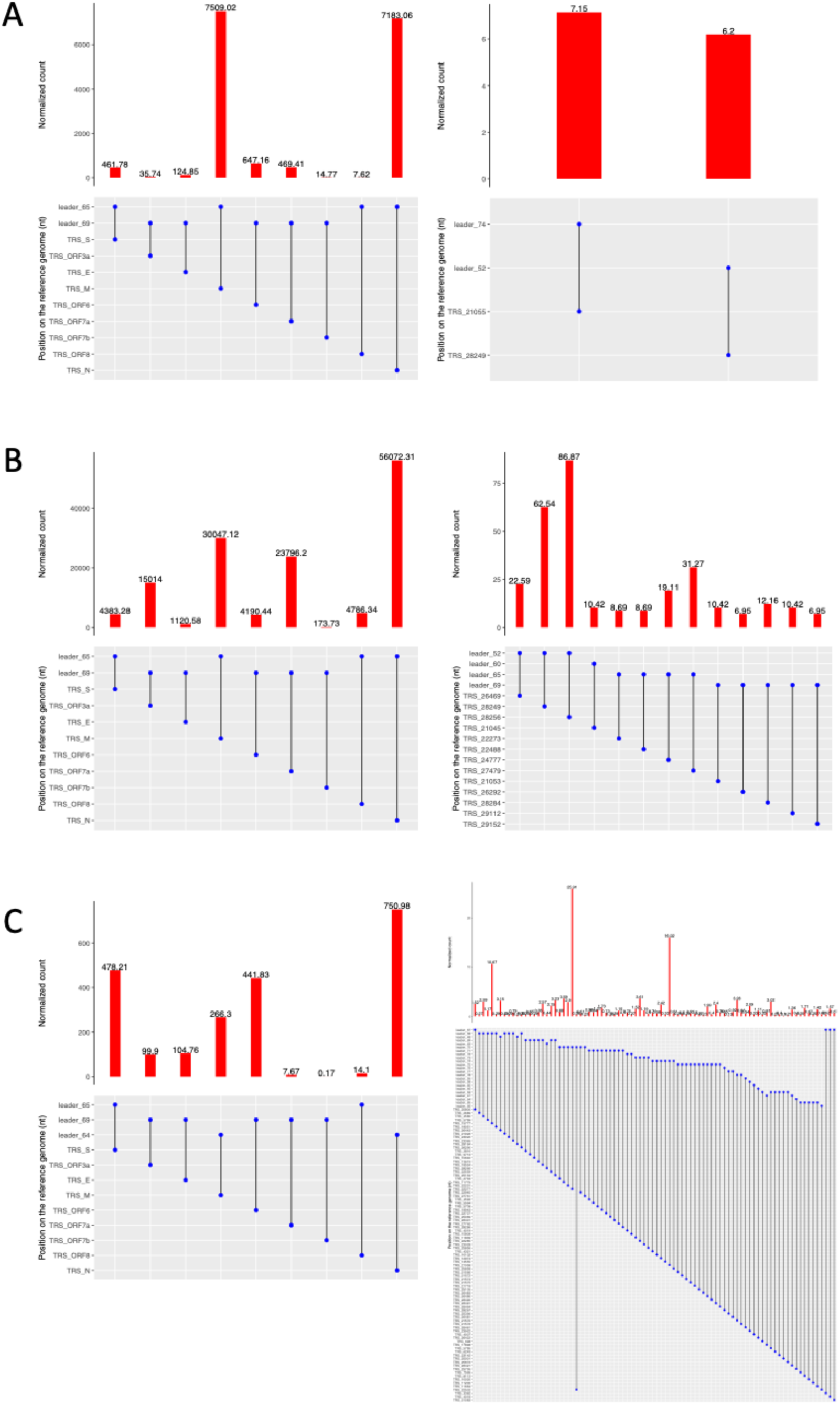
Analysis of leader TRS-gene junctions of reads with at least one primer sequence at either end in sequencing data from cell culture from (A) VeroE6 cells infected with SARS-CoV-2 (England/2/2020) and sequenced using an ARTIC Nanopore approach^16^ and (B) direct RNA sequencing of Cero CCL81 cells in culture infected with SARS-CoV-2 (National Culture Collection for Pathogens, Korea National Institute of Health, Korea)^2^. (C) Vero E6 cells were also infected with a near clinical isolate of known provenance and sequenced using the ARTIC Illumina approach.

### Evaluation of LeTRS on SARS-CoV-2 infection in cell culture

In order to assess the ability of LeTRS to identify the leader-TRS junctions from sequencing information, the tool was first evaluated on sequence data obtained from published SARS-CoV-2 infections in cell culture and our laboratory experiments conducted for this study. First, published data was used from sequencing viral RNA at 72 hrs post-infection in a cell culture model^16^. SARS-CoV-2 was sequenced using Nanopore from an amplicon-based approach (ARTIC)^16^ (Figure 2A, Table 1 and 2, Supplementary Tables 1 and 2). All of the major known leader-TRS gene junctions were identified. Interestingly, the nucleoprotein gene leader-TRS was approximately the same abundance as the membrane leader-TRS, whereas the other leader-TRSs were much lower. Two potential novel leader-TRS gene junctions were identified at positions 21,055 and 28,249 (Figure 2A, Table 2, Supplementary Table 2). The former is within the orf1b region and the latter within orf8. Second, data was analysed from a published experiment of cells infected (Figure 2B, Table 3 and 4, Supplementary Tables 3 and 4) and control sample (Tables 5 and 6) in culture using a direct RNA sequencing approach^2^. Analysis demonstrated a more expected pattern of abundance of the leader-TRS gene junctions (Figure 2B, Table 3 and Supplementary Table 3). The leader-TRS nucleoprotein gene junction was most abundant, and in general, the patten of abundance of the leader-TRS gene junctions for the major structural proteins followed the order of the gene junction along the genome. Novel low abundance leader-TRS gene junctions were also identified. One of these low abundance leader-TRSs gene junctions was also common to those found by the ARTIC amplicon analysis (Figure 2A and B, Table 2 and 4, Supplementary Table 2 and 4). Third, LeTRS was evaluated on sequencing data obtained from VeroE6 cells infected in culture with SARS-CoV-2 (SCV2-006). Here viral RNA, that had been prepared at 24 hrs post-infection, was amplified using the ARTIC approach and sequenced by Illumina (Figure 2C, Table 7 and 8, Supplementary Tables 5 and 6). As would be predicted the major leader-TRS gene junctions were identified, with the nucleoprotein one being the most abundant. Novel potential leader-gene junctions were also identified, including three that were greater in abundance than the other leader-gene junction. Some of the novel leader-TRS gene junctions from these three cell culture data sets shared the same first orf with the known sgmRNAs (Supplementary Tables 2, 4 and 6).

**Table 1.**
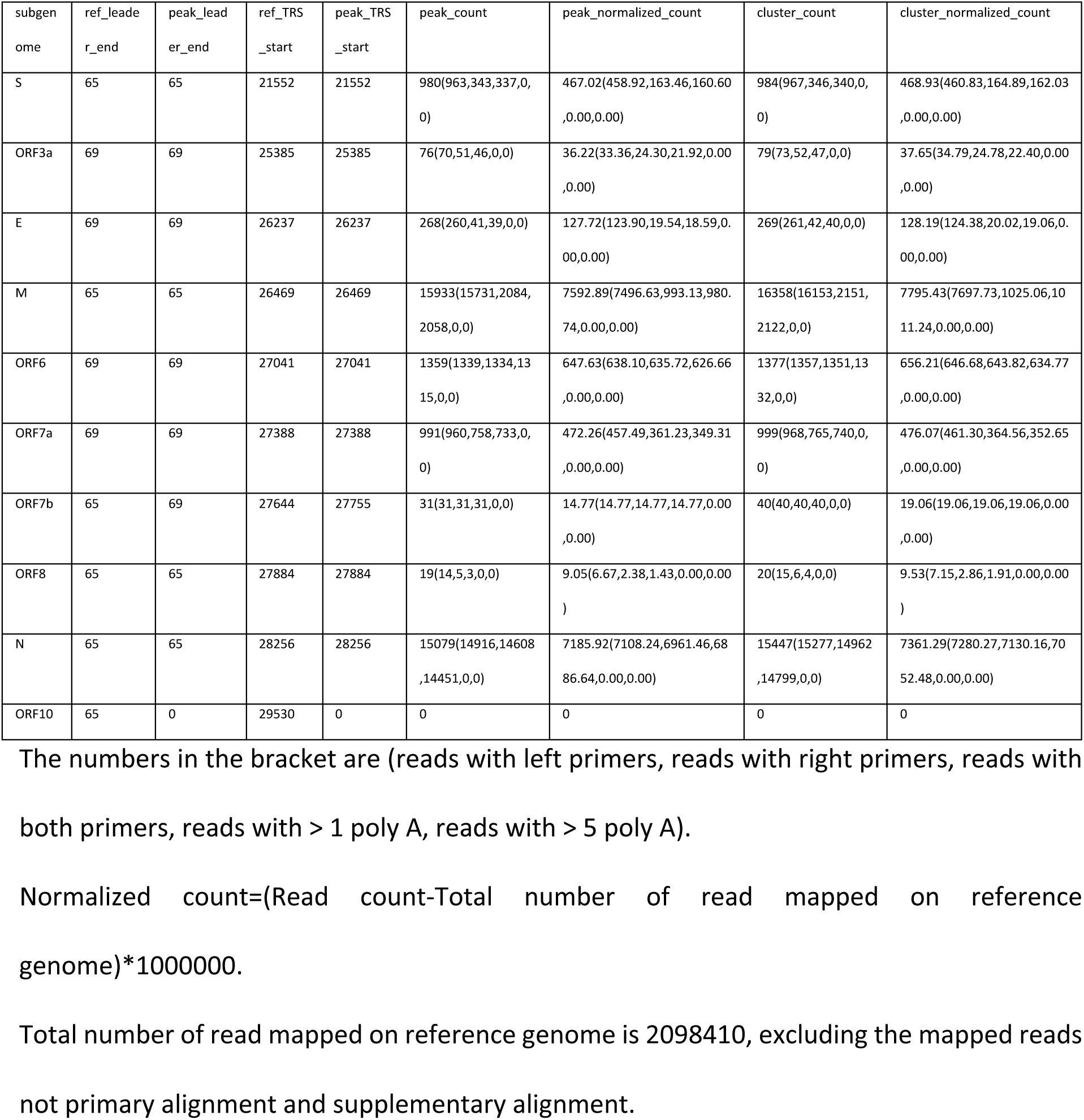
The LeTRS output table for known sgmRNA in the tested Nanopore ARTIC v3 primers amplicon sequencing data. “ref_leader_end” and “peak_leader_end” point to the reference position of the end of leader and the position of the end of leader identified in the most common reads (peak count) on the reference genome, and “ref_TRS_start” and “peak_TRS_start” refer to the reference position of the start of TRS and the position of the start of TRS identified in the most common reads (peak count) on the reference genome.

**Table 2.** The LeTRS output table for novel sgmRNA in the tested Nanopore ARTIC v3 primers amplicon sequencing data. “leader_end” and “TRS_start” refer to the position of the end of leader and the position of the start of TRS identified in the reads >10.

| subgenome | leader_end | TRS_start | nb_count | normalized_count |
| --- | --- | --- | --- | --- |
| 1 | 74 | 21055 | 15(13,15,13,0,0) | 7.15(6.20,7.15,6.20,0.00,0.00) |
| 2 | 52 | 28249 | 13(13,12,12,0,0) | 6.20(6.20,5.72,5.72,0.00,0.00) |

The numbers in the bracket are (reads with left primers, reads with right primers, reads with both primers, same junction on paired reads with at least a primer).
Normalized count=(Read count/Total number of read mapped on reference genome)\*1000000.
Total number of read mapped on reference genome is 2098410, excluding the mapped reads unpaired, not primary alignment and supplementary alignment.

**Table 3.**
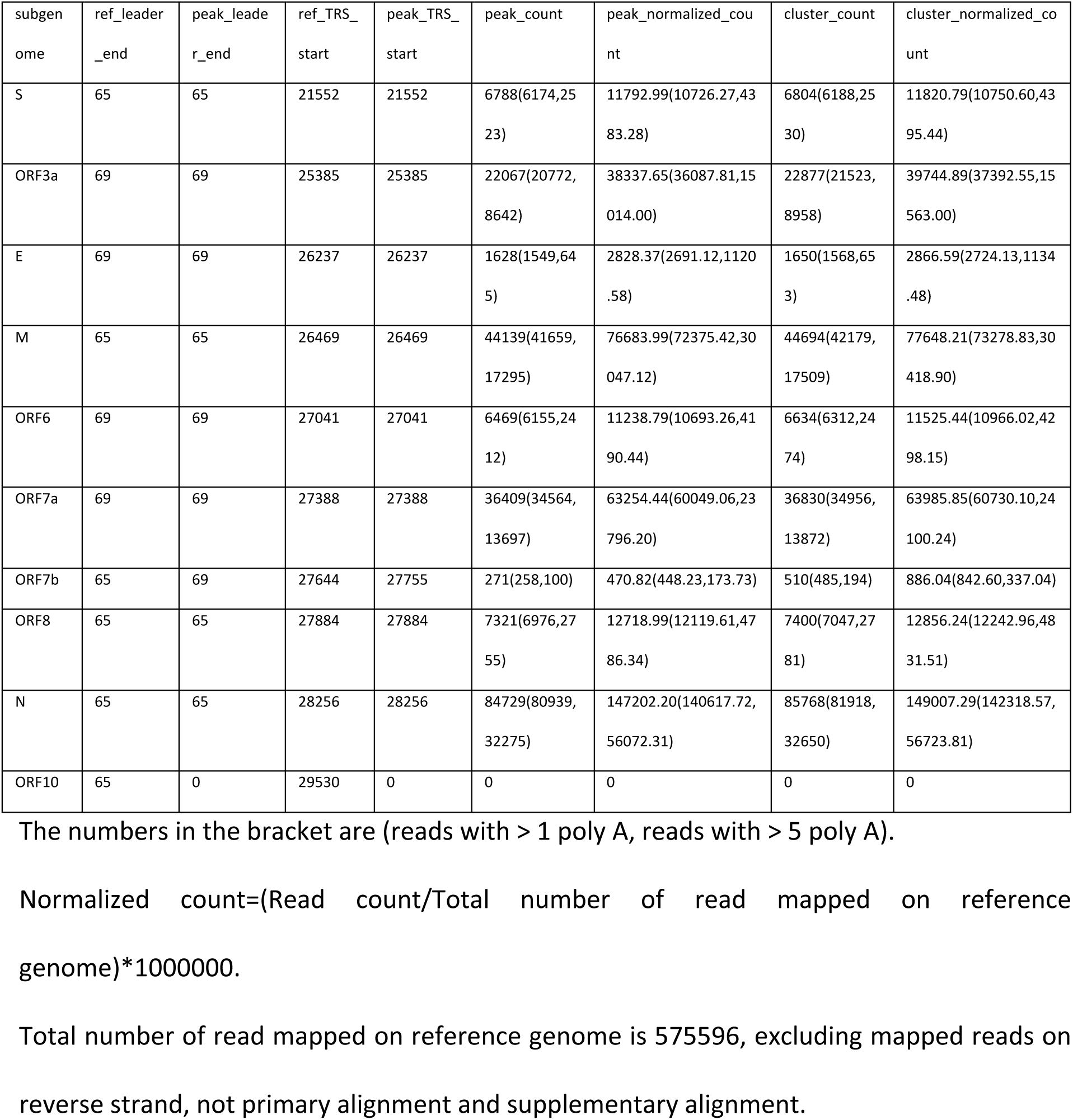
The LeTRS output table for known sgmRNA in the tested Nanopore direct RNA sequencing data. “ref_leader_end” and “peak_leader_end” point to the reference position of the end of leader and the position of the end of leader identified in the most common reads (peak count) on the reference genome, and “ref_TRS_start” and “peak_TRS_start” refer to the reference position of the start of TRS and the position of the start of TRS identified in the most common reads (peak count) on the reference genome.

**Table 5.**
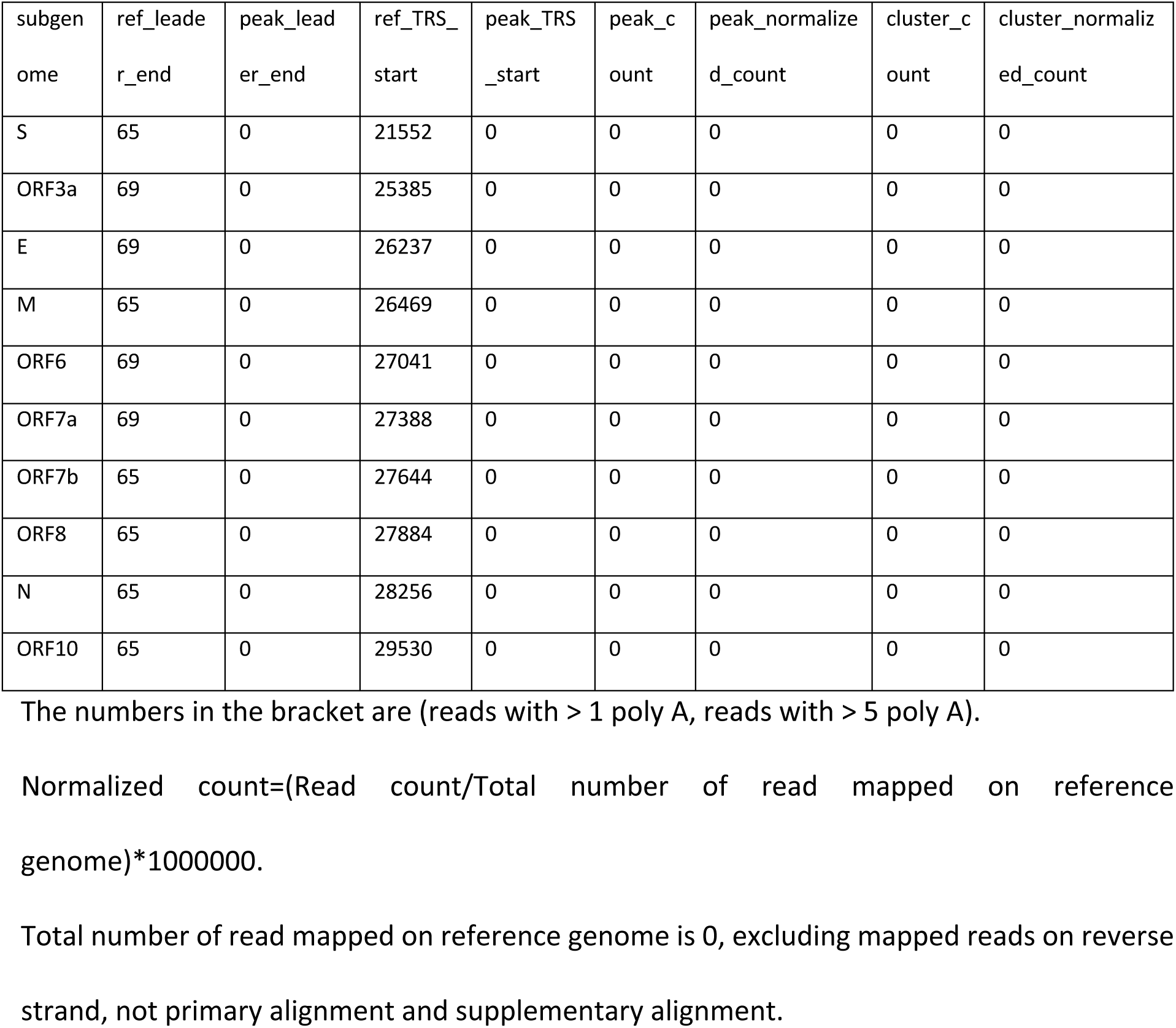
The LeTRS output table for known sgmRNA in the negative control of the tested nanopore direct RNA sequencing data. “ref_leader_end” and “peak_leader_end” point to the reference position of the end of leader and the position of the end of leader identified in the most common reads (peak count) on the reference genome, and “ref_TRS_start” and “peak_TRS_start” refer to the reference position of the start of TRS and the position of the start of TRS identified in the most common reads (peak count) on the reference genome.

| subgenome | ref_leader_end | peak_leader_end | ref_TRS_start | peak_TRS_start | peak_count | peak_normalized_count | cluster_count | cluster_normalized_count |
| --- | --- | --- | --- | --- | --- | --- | --- | --- |
| S | 65 | 0 | 21552 | 0 | 0 | 0 | 0 | 0 |
| ORF3a | 69 | 0 | 25385 | 0 | 0 | 0 | 0 | 0 |
| E | 69 | 0 | 26237 | 0 | 0 | 0 | 0 | 0 |
| M | 65 | 0 | 26469 | 0 | 0 | 0 | 0 | 0 |
| ORF6 | 69 | 0 | 27041 | 0 | 0 | 0 | 0 | 0 |
| ORF7a | 69 | 0 | 27388 | 0 | 0 | 0 | 0 | 0 |
| ORF7b | 65 | 0 | 27644 | 0 | 0 | 0 | 0 | 0 |
| ORF8 | 65 | 0 | 27884 | 0 | 0 | 0 | 0 | 0 |
| N | 65 | 0 | 28256 | 0 | 0 | 0 | 0 | 0 |
| ORF10 | 65 | 0 | 29530 | 0 | 0 | 0 | 0 | 0 |

The numbers in the bracket are (reads with > 1 poly A, reads with > 5 poly A).
Normalized count=(Read count/Total number of read mapped on reference genome)\*1000000.
Total number of read mapped on reference genome is 0, excluding mapped reads on reverse strand, not primary alignment and supplementary alignment.

**Table 6.**
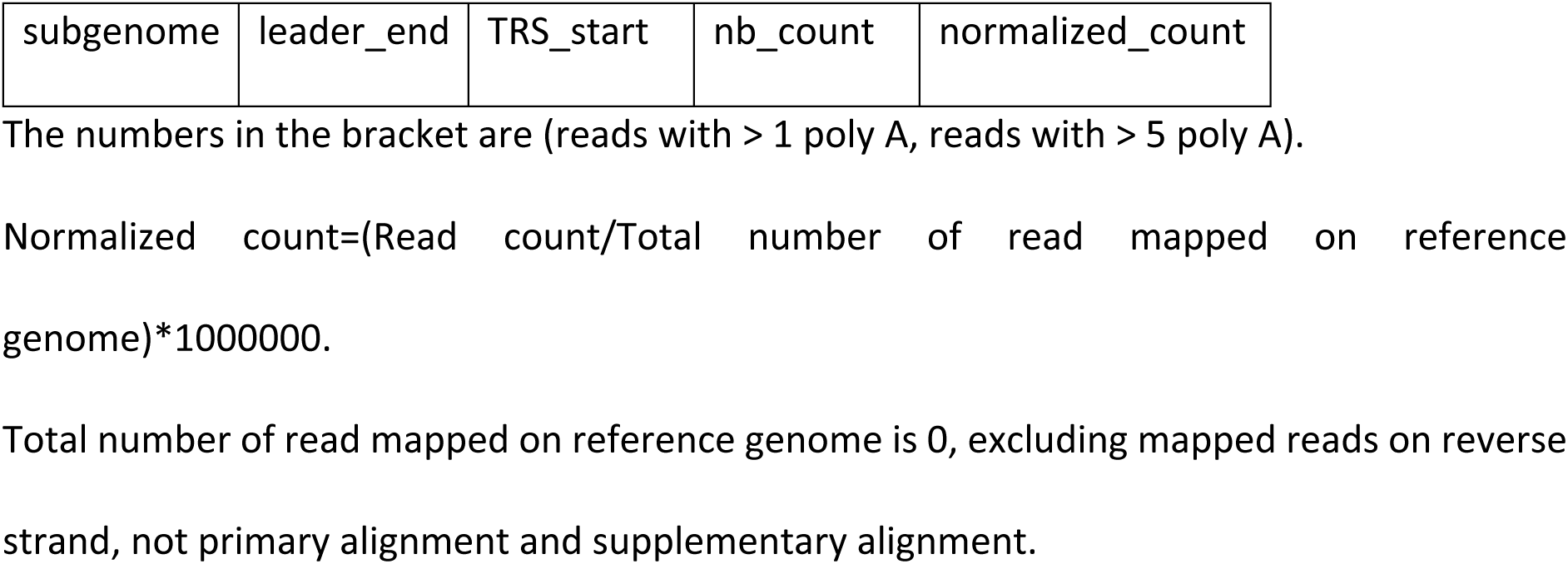
The LeTRS output table for novel sgmRNA in the negative control of the tested nanopore direct RNA sequencing data. “leader_end” and “TRS_start” refer to the position of the end of leader and the position of the start of TRS identified in the reads >10.

**Table 7.**
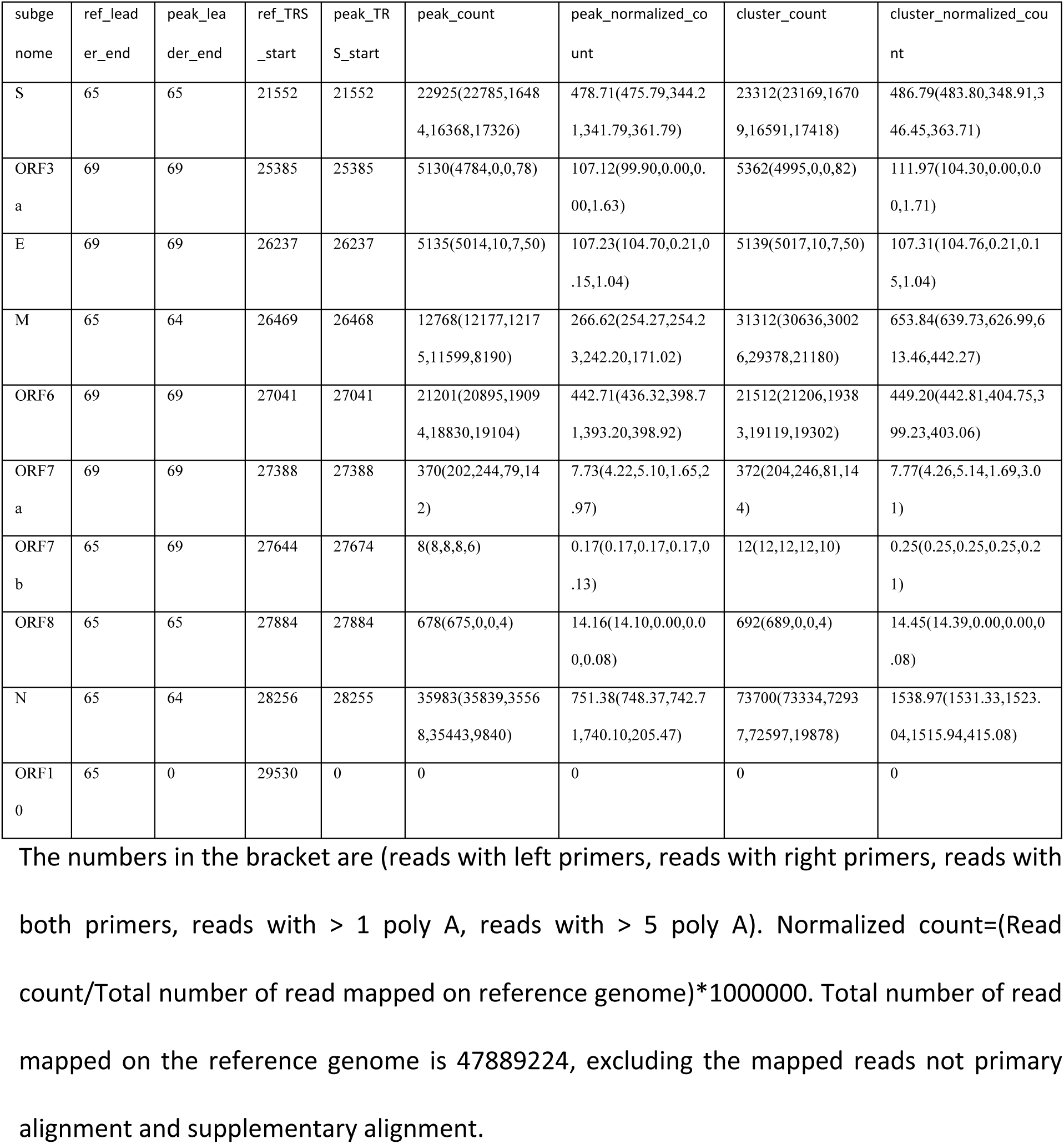
The LeTRS output table for known sgmRNA in the tested Illumina ARTIC v3 primers amplicon sequencing data. “ref_leader_end” and “peak_leader_end” point to the reference position of the end of leader and the position of the end of leader identified in the most common reads (peak count) on the reference genome, and “ref_TRS_start” and “peak_TRS_start” refer to the reference position of the start of TRS and the position of the start of TRS identified in the most common reads (peak count) on the reference genome.

| subgenome | ref_leader_end | peak_leader_end | ref_TRS_start | peak_TRS_start | peak_count | peak_normalized_count | cluster_count | cluster_normalized_count |
| --- | --- | --- | --- | --- | --- | --- | --- | --- |
| S | 65 | 65 | 21552 | 21552 | 22925(22785,1648<br>4,16368,17326) | 478.71(475.79,344.2<br>1,341.79,361.79) | 23312(23169,1670<br>9,16591,17418) | 486.79(483.80,348.91,3<br>46.45,363.71) |
| ORF3a | 69 | 69 | 25385 | 25385 | 5130(4784,0,0,78) | 107.12(99.90,0.00,0.<br>00,1.63) | 5362(4995,0,0,82) | 111.97(104.30,0.00,0.0<br>0,1.71) |
| E | 69 | 69 | 26237 | 26237 | 5135(5014,10,7,50) | 107.23(104.70,0.21,0<br>.15,1.04) | 5139(5017,10,7,50) | 107.31(104.76,0.21,0.1<br>5,1.04) |
| M | 65 | 64 | 26469 | 26468 | 12768(12177,1217<br>5,11599,8190) | 266.62(254.27,254.2<br>3,242.20,171.02) | 31312(30636,3002<br>6,29378,21180) | 653.84(639.73,626.99,6<br>13.46,442.27) |
| ORF6 | 69 | 69 | 27041 | 27041 | 21201(20895,1909<br>4,18830,19104) | 442.71(436.32,398.7<br>1,393.20,398.92) | 21512(21206,1938<br>3,19119,19302) | 449.20(442.81,404.75,3<br>99.23,403.06) |
| ORF7a | 69 | 69 | 27388 | 27388 | 370(202,244,79,14<br>2) | 7.73(4.22,5.10,1.65,2<br>.97) | 372(204,246,81,14<br>4) | 7.77(4.26,5.14,1.69,3.0<br>1) |
| ORF7b | 65 | 69 | 27644 | 27674 | 8(8,8,8,6) | 0.17(0.17,0.17,0.17,0<br>.13) | 12(12,12,12,10) | 0.25(0.25,0.25,0.25,0.2<br>1) |
| ORF8 | 65 | 65 | 27884 | 27884 | 678(675,0,0,4) | 14.16(14.10,0.00,0.0<br>0,0.08) | 692(689,0,0,4) | 14.45(14.39,0.00,0.00,0<br>.08) |
| N | 65 | 64 | 28256 | 28255 | 35983(35839,3556<br>8,35443,9840) | 751.38(748.37,742.7<br>1,740.10,205.47) | 73700(73334,7293<br>7,72597,19878) | 1538.97(1531.33,1523.<br>04,1515.94,415.08) |
| ORF10 | 65 | 0 | 29530 | 0 | 0 | 0 | 0 | 0 |

The numbers in the bracket are (reads with left primers, reads with right primers, reads with both primers, reads with > 1 poly A, reads with > 5 poly A). Normalized count=(Read count/Total number of read mapped on reference genome)\*1000000. Total number of read mapped on the reference genome is 47889224, excluding the mapped reads not primary alignment and supplementary alignment.

**Table 8.**
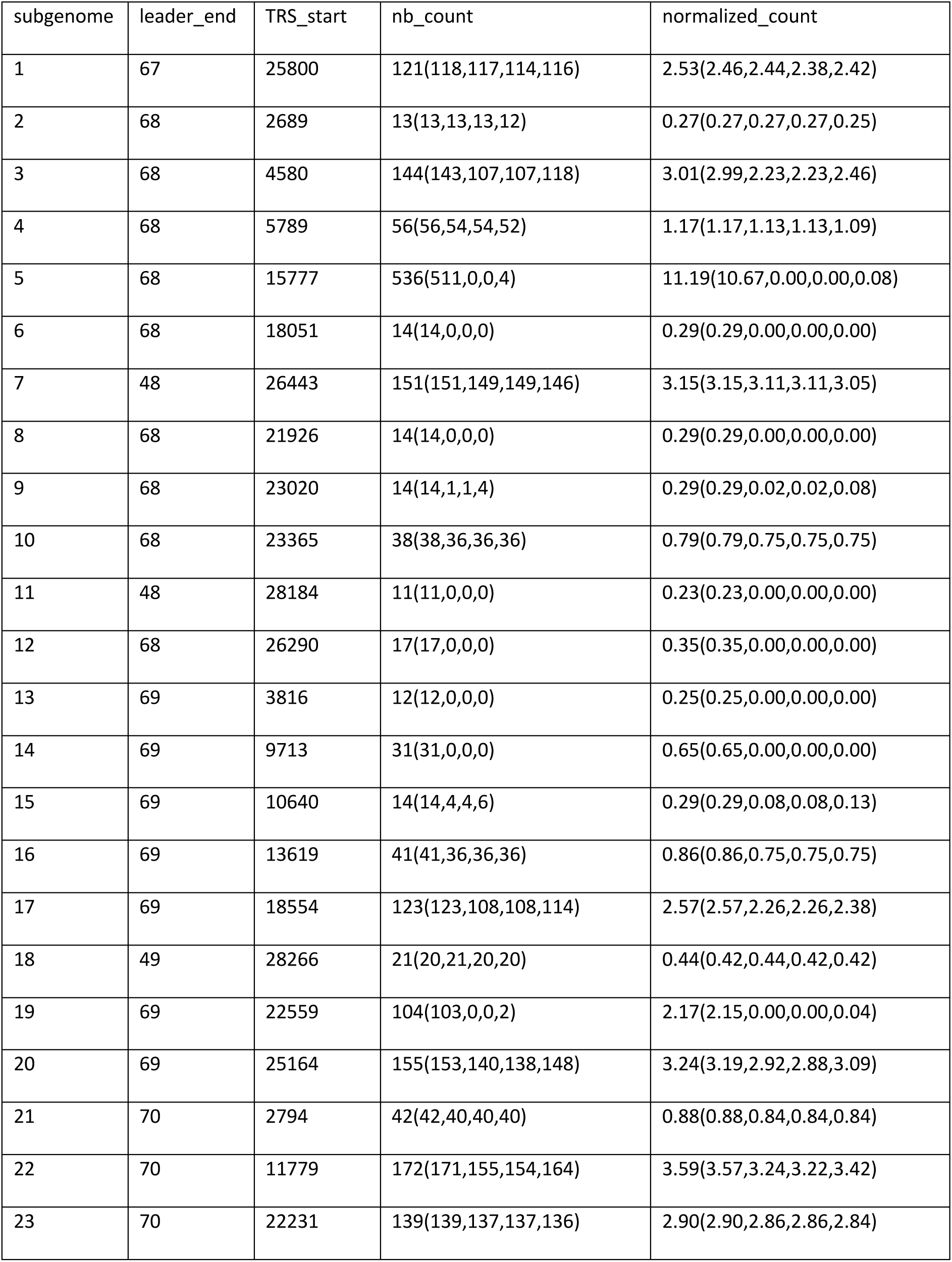
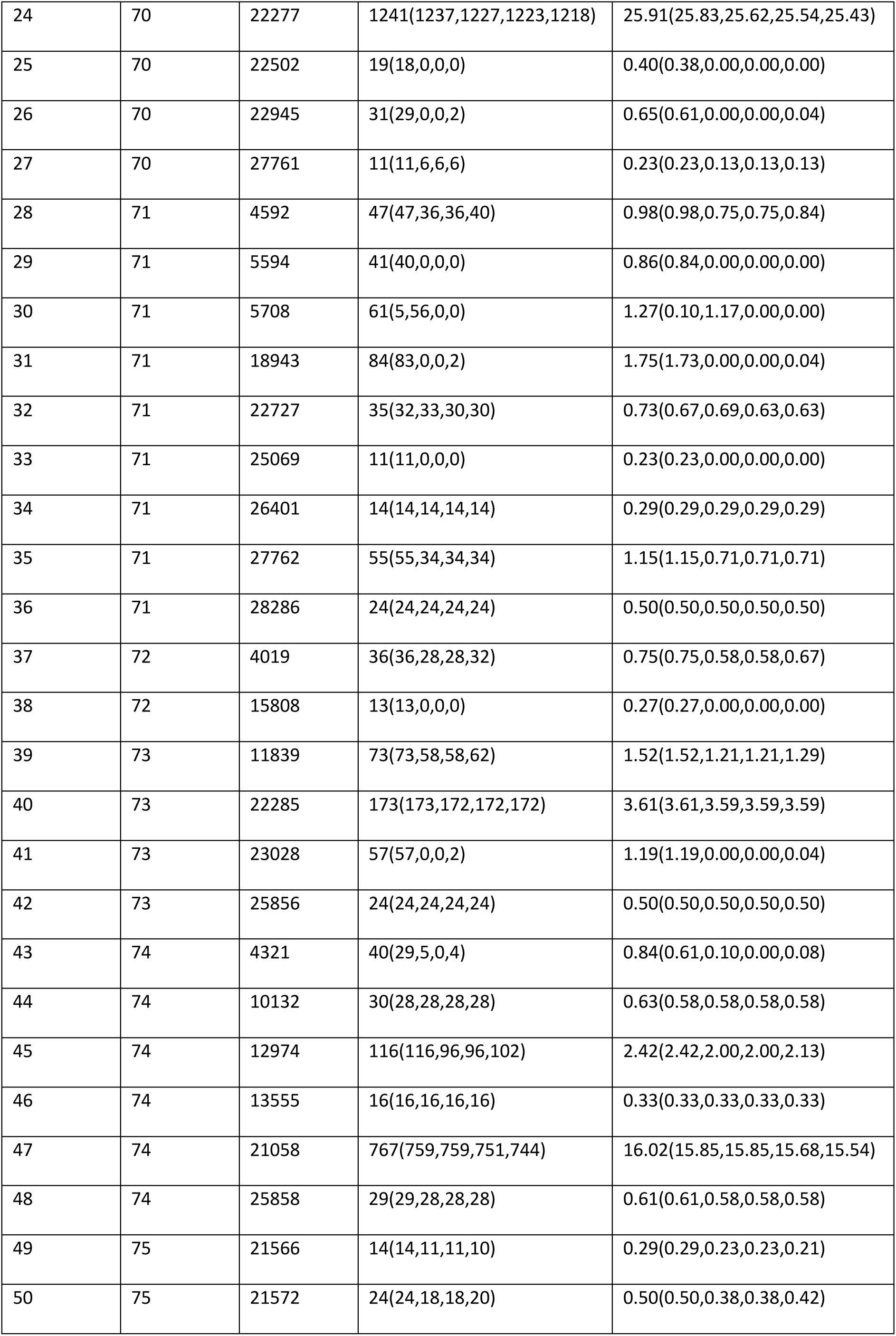
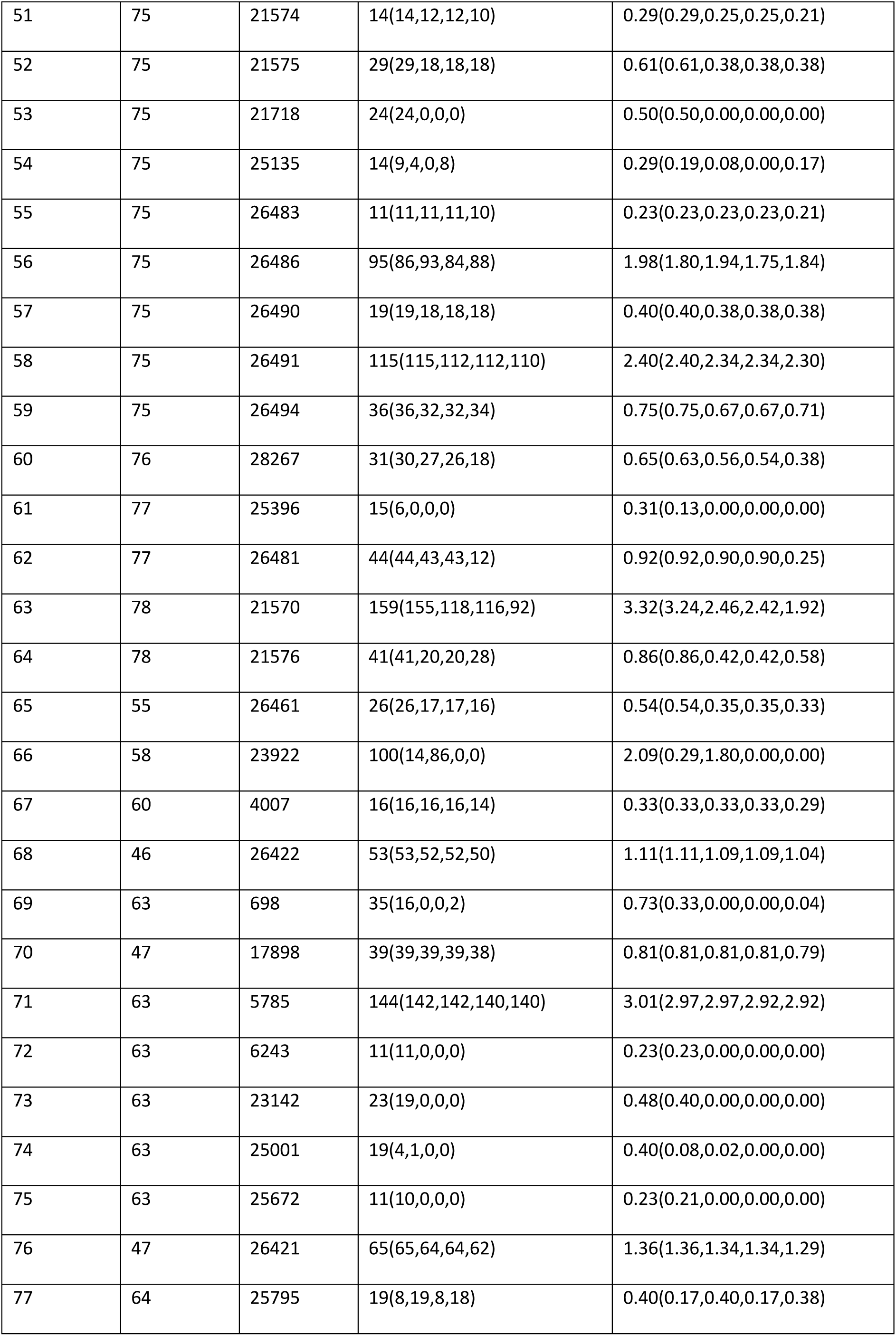
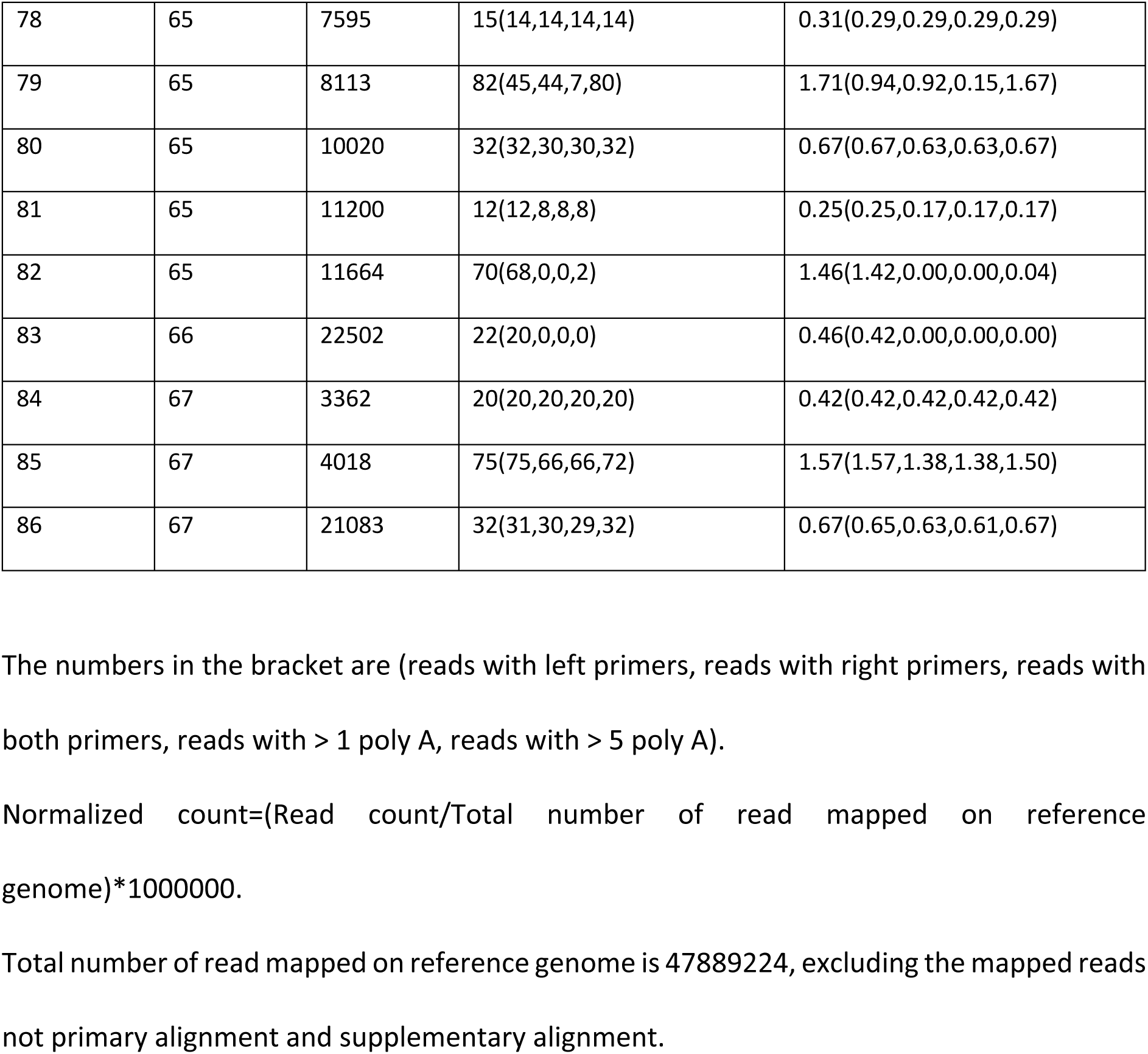
The LeTRS output table for novel sgmRNA in the tested Nanopore ARTIC v3 primers amplicon sequencing data. “leader_end” and “TRS_start” refer to the position of the end of leader and the position of the start of TRS identified in the reads >10.

| subgenome | leader_end | TRS_start | nb_count | normalized_count |
| --- | --- | --- | --- | --- |
| 1 | 67 | 25800 | 121(118,117,114,116) | 2.53(2.46,2.44,2.38,2.42) |
| 2 | 68 | 2689 | 13(13,13,13,12) | 0.27(0.27,0.27,0.27,0.25) |
| 3 | 68 | 4580 | 144(143,107,107,118) | 3.01(2.99,2.23,2.23,2.46) |
| 4 | 68 | 5789 | 56(56,54,54,52) | 1.17(1.17,1.13,1.13,1.09) |
| 5 | 68 | 15777 | 536(511,0,0,4) | 11.19(10.67,0.00,0.00,0.08) |
| 6 | 68 | 18051 | 14(14,0,0,0) | 0.29(0.29,0.00,0.00,0.00) |
| 7 | 48 | 26443 | 151(151,149,149,146) | 3.15(3.15,3.11,3.11,3.05) |
| 8 | 68 | 21926 | 14(14,0,0,0) | 0.29(0.29,0.00,0.00,0.00) |
| 9 | 68 | 23020 | 14(14,1,1,4) | 0.29(0.29,0.02,0.02,0.08) |
| 10 | 68 | 23365 | 38(38,36,36,36) | 0.79(0.79,0.75,0.75,0.75) |
| 11 | 48 | 28184 | 11(11,0,0,0) | 0.23(0.23,0.00,0.00,0.00) |
| 12 | 68 | 26290 | 17(17,0,0,0) | 0.35(0.35,0.00,0.00,0.00) |
| 13 | 69 | 3816 | 12(12,0,0,0) | 0.25(0.25,0.00,0.00,0.00) |
| 14 | 69 | 9713 | 31(31,0,0,0) | 0.65(0.65,0.00,0.00,0.00) |
| 15 | 69 | 10640 | 14(14,4,4,6) | 0.29(0.29,0.08,0.08,0.13) |
| 16 | 69 | 13619 | 41(41,36,36,36) | 0.86(0.86,0.75,0.75,0.75) |
| 17 | 69 | 18554 | 123(123,108,108,114) | 2.57(2.57,2.26,2.26,2.38) |
| 18 | 49 | 28266 | 21(20,21,20,20) | 0.44(0.42,0.44,0.42,0.42) |
| 19 | 69 | 22559 | 104(103,0,0,2) | 2.17(2.15,0.00,0.00,0.04) |
| 20 | 69 | 25164 | 155(153,140,138,148) | 3.24(3.19,2.92,2.88,3.09) |
| 21 | 70 | 2794 | 42(42,40,40,40) | 0.88(0.88,0.84,0.84,0.84) |
| 22 | 70 | 11779 | 172(171,155,154,164) | 3.59(3.57,3.24,3.22,3.42) |
| 23 | 70 | 22231 | 139(139,137,137,136) | 2.90(2.90,2.86,2.86,2.84) |
| 24 | 70 | 22277 | 1241(1237,1227,1223,1218) | 25.91(25.83,25.62,25.54,25.43) |
| 25 | 70 | 22502 | 19(18,0,0,0) | 0.40(0.38,0.00,0.00,0.00) |
| 26 | 70 | 22945 | 31(29,0,0,2) | 0.65(0.61,0.00,0.00,0.04) |
| 27 | 70 | 27761 | 11(11,6,6,6) | 0.23(0.23,0.13,0.13,0.13) |
| 28 | 71 | 4592 | 47(47,36,36,40) | 0.98(0.98,0.75,0.75,0.84) |
| 29 | 71 | 5594 | 41(40,0,0,0) | 0.86(0.84,0.00,0.00,0.00) |
| 30 | 71 | 5708 | 61(5,56,0,0) | 1.27(0.10,1.17,0.00,0.00) |
| 31 | 71 | 18943 | 84(83,0,0,2) | 1.75(1.73,0.00,0.00,0.04) |
| 32 | 71 | 22727 | 35(32,33,30,30) | 0.73(0.67,0.69,0.63,0.63) |
| 33 | 71 | 25069 | 11(11,0,0,0) | 0.23(0.23,0.00,0.00,0.00) |
| 34 | 71 | 26401 | 14(14,14,14,14) | 0.29(0.29,0.29,0.29,0.29) |
| 35 | 71 | 27762 | 55(55,34,34,34) | 1.15(1.15,0.71,0.71,0.71) |
| 36 | 71 | 28286 | 24(24,24,24,24) | 0.50(0.50,0.50,0.50,0.50) |
| 37 | 72 | 4019 | 36(36,28,28,32) | 0.75(0.75,0.58,0.58,0.67) |
| 38 | 72 | 15808 | 13(13,0,0,0) | 0.27(0.27,0.00,0.00,0.00) |
| 39 | 73 | 11839 | 73(73,58,58,62) | 1.52(1.52,1.21,1.21,1.29) |
| 40 | 73 | 22285 | 173(173,172,172,172) | 3.61(3.61,3.59,3.59,3.59) |
| 41 | 73 | 23028 | 57(57,0,0,2) | 1.19(1.19,0.00,0.00,0.04) |
| 42 | 73 | 25856 | 24(24,24,24,24) | 0.50(0.50,0.50,0.50,0.50) |
| 43 | 74 | 4321 | 40(29,5,0,4) | 0.84(0.61,0.10,0.00,0.08) |
| 44 | 74 | 10132 | 30(28,28,28,28) | 0.63(0.58,0.58,0.58,0.58) |
| 45 | 74 | 12974 | 116(116,96,96,102) | 2.42(2.42,2.00,2.00,2.13) |
| 46 | 74 | 13555 | 16(16,16,16,16) | 0.33(0.33,0.33,0.33,0.33) |
| 47 | 74 | 21058 | 767(759,759,751,744) | 16.02(15.85,15.85,15.68,15.54) |
| 48 | 74 | 25858 | 29(29,28,28,28) | 0.61(0.61,0.58,0.58,0.58) |
| 49 | 75 | 21566 | 14(14,11,11,10) | 0.29(0.29,0.23,0.23,0.21) |
| 50 | 75 | 21572 | 24(24,18,18,20) | 0.50(0.50,0.38,0.38,0.42) |
| 51 | 75 | 21574 | 14(14,12,12,10) | 0.29(0.29,0.25,0.25,0.21) |
| 52 | 75 | 21575 | 29(29,18,18,18) | 0.61(0.61,0.38,0.38,0.38) |
| 53 | 75 | 21718 | 24(24,0,0,0) | 0.50(0.50,0.00,0.00,0.00) |
| 54 | 75 | 25135 | 14(9,4,0,8) | 0.29(0.19,0.08,0.00,0.17) |
| 55 | 75 | 26483 | 11(11,11,11,10) | 0.23(0.23,0.23,0.23,0.21) |
| 56 | 75 | 26486 | 95(86,93,84,88) | 1.98(1.80,1.94,1.75,1.84) |
| 57 | 75 | 26490 | 19(19,18,18,18) | 0.40(0.40,0.38,0.38,0.38) |
| 58 | 75 | 26491 | 115(115,112,112,110) | 2.40(2.40,2.34,2.34,2.30) |
| 59 | 75 | 26494 | 36(36,32,32,34) | 0.75(0.75,0.67,0.67,0.71) |
| 60 | 76 | 28267 | 31(30,27,26,18) | 0.65(0.63,0.56,0.54,0.38) |
| 61 | 77 | 25396 | 15(6,0,0,0) | 0.31(0.13,0.00,0.00,0.00) |
| 62 | 77 | 26481 | 44(44,43,43,12) | 0.92(0.92,0.90,0.90,0.25) |
| 63 | 78 | 21570 | 159(155,118,116,92) | 3.32(3.24,2.46,2.42,1.92) |
| 64 | 78 | 21576 | 41(41,20,20,28) | 0.86(0.86,0.42,0.42,0.58) |
| 65 | 55 | 26461 | 26(26,17,17,16) | 0.54(0.54,0.35,0.35,0.33) |
| 66 | 58 | 23922 | 100(14,86,0,0) | 2.09(0.29,1.80,0.00,0.00) |
| 67 | 60 | 4007 | 16(16,16,16,14) | 0.33(0.33,0.33,0.33,0.29) |
| 68 | 46 | 26422 | 53(53,52,52,50) | 1.11(1.11,1.09,1.09,1.04) |
| 69 | 63 | 698 | 35(16,0,0,2) | 0.73(0.33,0.00,0.00,0.04) |
| 70 | 47 | 17898 | 39(39,39,39,38) | 0.81(0.81,0.81,0.81,0.79) |
| 71 | 63 | 5785 | 144(142,142,140,140) | 3.01(2.97,2.97,2.92,2.92) |
| 72 | 63 | 6243 | 11(11,0,0,0) | 0.23(0.23,0.00,0.00,0.00) |
| 73 | 63 | 23142 | 23(19,0,0,0) | 0.48(0.40,0.00,0.00,0.00) |
| 74 | 63 | 25001 | 19(4,1,0,0) | 0.40(0.08,0.02,0.00,0.00) |
| 75 | 63 | 25672 | 11(10,0,0,0) | 0.23(0.21,0.00,0.00,0.00) |
| 76 | 47 | 26421 | 65(65,64,64,62) | 1.36(1.36,1.34,1.34,1.29) |
| 77 | 64 | 25795 | 19(8,19,8,18) | 0.40(0.17,0.40,0.17,0.38) |
| 78 | 65 | 7595 | 15(14,14,14,14) | 0.31(0.29,0.29,0.29,0.29) |
| 79 | 65 | 8113 | 82(45,44,7,80) | 1.71(0.94,0.92,0.15,1.67) |
| 80 | 65 | 10020 | 32(32,30,30,32) | 0.67(0.67,0.63,0.63,0.67) |
| 81 | 65 | 11200 | 12(12,8,8,8) | 0.25(0.25,0.17,0.17,0.17) |
| 82 | 65 | 11664 | 70(68,0,0,2) | 1.46(1.42,0.00,0.00,0.04) |
| 83 | 66 | 22502 | 22(20,0,0,0) | 0.46(0.42,0.00,0.00,0.00) |
| 84 | 67 | 3362 | 20(20,20,20,20) | 0.42(0.42,0.42,0.42,0.42) |
| 85 | 67 | 4018 | 75(75,66,66,72) | 1.57(1.57,1.38,1.38,1.50) |
| 86 | 67 | 21083 | 32(31,30,29,32) | 0.67(0.65,0.63,0.61,0.67) |

The numbers in the bracket are (reads with left primers, reads with right primers, reads with both primers, reads with > 1 poly A, reads with > 5 poly A).
Normalized count=(Read count/Total number of read mapped on reference genome)\*1000000.
Total number of read mapped on reference genome is 47889224, excluding the mapped reads not primary alignment and supplementary alignment.

### Comparison with other informatic tools that can identify leader TRS gene junctions

Periscope v0.08a is another tool that was developed to identify sgmRNA from Illumina and Nanopore ARTIC amplicon sequencing data^17^. The tool functions based on searching a 32 nt leader sequence (genomic position: 34-65) and anchoring the known TRS-orf boundaries on the reads for identification of known sgmRNAs. Periscope does not take into consideration the sequences and distance between the leader and TRS-orf boundaries. Periscope can analyse ARTIC amplicon sequencing data, whereas LeTRS can also input a variety of different amplicon and direct RNA sequencing data. Given the very large sequencing datasets being generated as part of the global effort to sequence SARS-CoV-2 the performance of LeTRS was compared to Periscope in terms of computation time. Illumina sequencing data from a nasopharyngeal sample of a human patient with COVID-19 and Nanopore ARTIC amplicon published cell culture data sets were used for comparison. This used the number of reads with at least one primer sequence at either end in the LeTRS and the number of “High Quality” reads (the reads with both 32 nts leader sequences and known TRS-orf boundary) in Periscope. Periscope was run with the default setting^17^. Both LeTRS and Periscope identified a similar number of reads for both data sets (Supplementary Figure 2, Tables 1 and 3 and Supplementary table 7). With 16 CPU cores, the run times for LeTRS was 1m 40.692s and 2m 14.911s for these tested Illumina and Nanopore data sets, respectively, and for Periscope these were 7m 31.183s and 16m 49.448s. We also tested the data from ARTIC Illumina cell culture data, but Periscope had an error.

### Analysis of sequencing data from longitudinal nasopharyngeal samples taken from two non-human primate models of COVID-19 indicated multi-phasic sgmRNA synthesis

Part of the difficulty of studying SARS-CoV-2 and the disease COVID-19 is establishing the sequence of events from the start of infection. Most samples from humans are from nasopharyngeal aspirates taken when clinical symptoms develop. This tends to be 5 to 6 days post-exposure. In the absence of a human challenge model, animal models can be used to study the kinetics of SARS-CoV-2^18, 19^. Two separate non-human primate models, cynomolgus and rhesus macaques, were established for the study of SARS-CoV-2 that mirrored disease in the majority of humans^18^. To study the pattern of sgmRNA synthesis over the course of infection, nasopharyngeal samples were sequentially gathered daily from one day post-infection up to 18 days post-infection from the two NHP models. RNA was purified from these longitudinal samples as well as the inoculum virus and viral RNA sequenced using the ARTIC approach on the Illumina platform.

Analysis of the sequence data from the inoculum used to infect the NHPs indicated that leader gene junctions could be identified (Supplementary Figure 3, Supplementary_Table_8), but these did not follow the pattern of abundance of leader TRS-gene junctions found in infected cells in culture, where the leader TRS-N gene junction was most abundant (Figure 1C). In contrast, analysis of the longitudinal sequencing data from nasopharyngeal aspirates from the non-human primate model identified leader TRS-gene junctions associated with the major sgmRNAs (Figure 3, Supplementary_Table_9) as well as novel leader-TRS gene junction sites (Supplementary Figures 4 and 5). Analysing the abundance of the leader TRS-gene junctions for both model species over the course of infection revealed a phasic nature of sgmRNA synthesis. The leader TRS nucleoprotein gene junction was the most abundant, and there was a similar phasic pattern of potential sgmRNA synthesis with Illumina ARTIC method (Figure 3). For both species, viral load and hence sgmRNA synthesis had dropped by Day 8 and Day 9.

**Figure 3.**
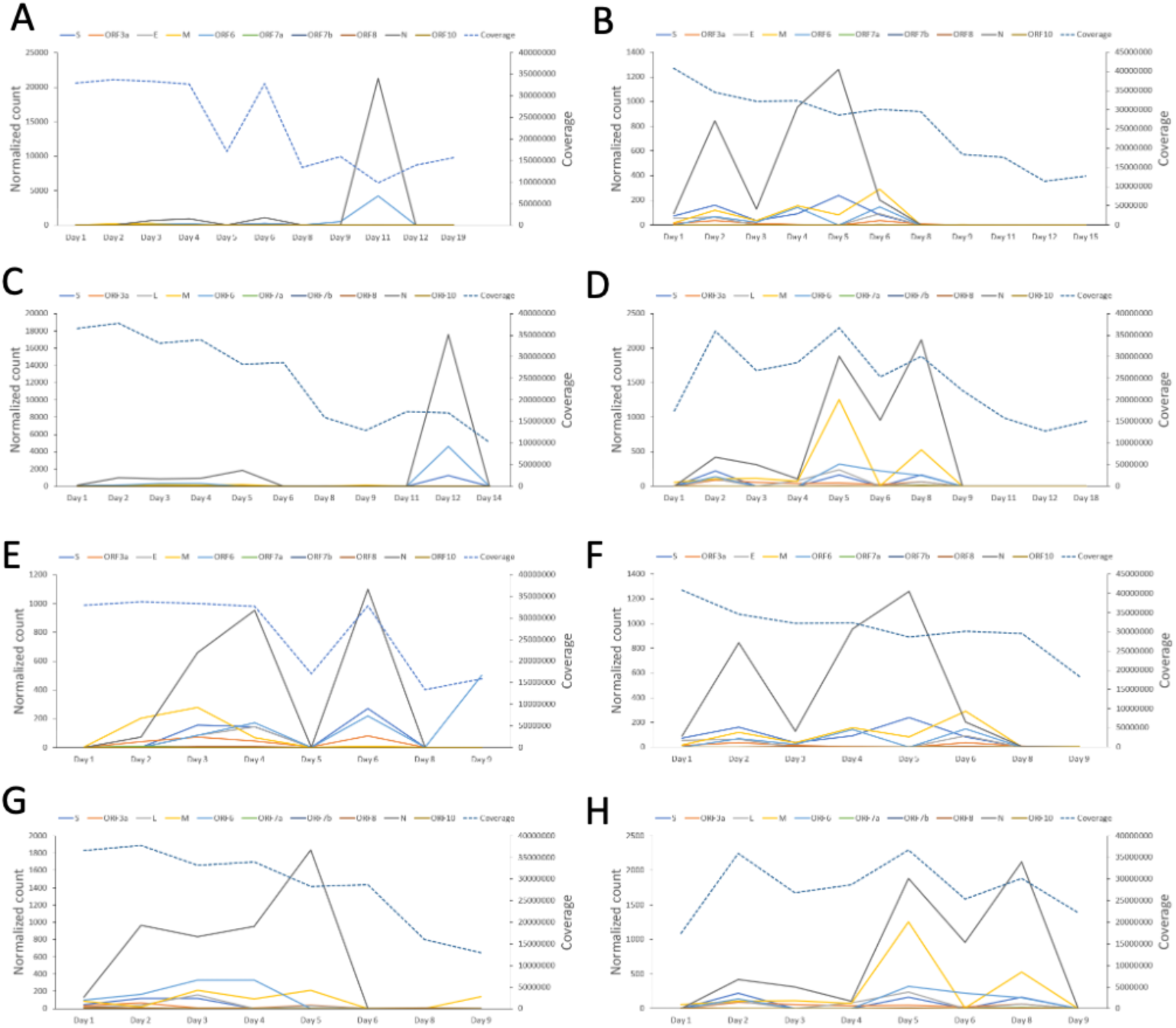
Analysis of leader TRS-gene junction abundance of reads with at least one primer sequence at either end in longitudinal nasopharyngeal samples (daily indicated on the x-axis) taken from two non-human primate models of SARS-CoV-2 in groups. The normalised count (Read count/total number of reads mapped on the reference genome)*1,000,000) of the leader TRS-gene junction abundance is shown on the left-hand Y-axis with each unique junction colour coded. The right-hand Y axis is a measure of the total depth of coverage for SARS-CoV-2 in that sample. Note the two scales are different. SARS-CoV-2 was amplified using the ARTIC approach and sequenced by Illumina. The data is organised into groups of animals for the cynomolgus macaque groups 1 and 2 (A/E and B/F), and rhesus macaque groups 1 and 2 (C/G and D/H). E, F, G and H zoom in to see the details of A, B, C and D for Day1 to Day9. The data correspond to Supplementary Table 9.

### Analysis of leader TRS-gene junction in human samples revealed expected and aberrant abundances

To investigate the pattern of leader-TRS gene junction abundance during infection of SARS-CoV-2 in humans, nasopharyngeal swabs form patients with COVID-19 were sequenced by the ARTIC approach using either Illumina (as part of COG-UK) (N=15 patients) (Figure 4, Supplementary Table 7) or by Oxford Nanopore (as part of ISARIC-4C) (N=15 patients) (Figure 5, Supplementary Table 10). In a number of samples, leader-TRS gene junctions were identified, and followed an expected pattern, with the nucleoprotein gene junction being the most abundant (e.g., Sample 1 in Figures 4A and B, Patient 2 day1 in Figure 5A and B). However, in several of the samples there was very large representation of single leader-TRS gene junction (e.g., Sample 4 and 5 in Figures 4A and B). These tended to map to the nucleoprotein gene (Sample 5, 8 and 13 Figures 4A and B). The heterogeneity in abundance of leader gene junctions was reminiscent for that from the non-human primate study with a defined and expected pattern near the start of infection but then becoming phasic. The samples gathered under ISARIC-4C were from hospitalised patients and permitted analysis in relation to reported date of symptom onset and sequential sampling. In general, the data indicated that on first sample on admission to hospital contained an abundance of leader-TRS gene junctions which resembled the pattern seen in infected cells (Patient 6 day1 and day9 in Figures 5A and B). However, with further days post-sample, e.g. (Patient 7 day7 Figures 5A and B), the leader-TRS N gene junction was the most abundant and far exceeded any other detectable species. The abundance of leader-TRS N gene junction in the patients at a later stage of infection followed that observed in the NHP model (Figure 3).

**Figure 4.**
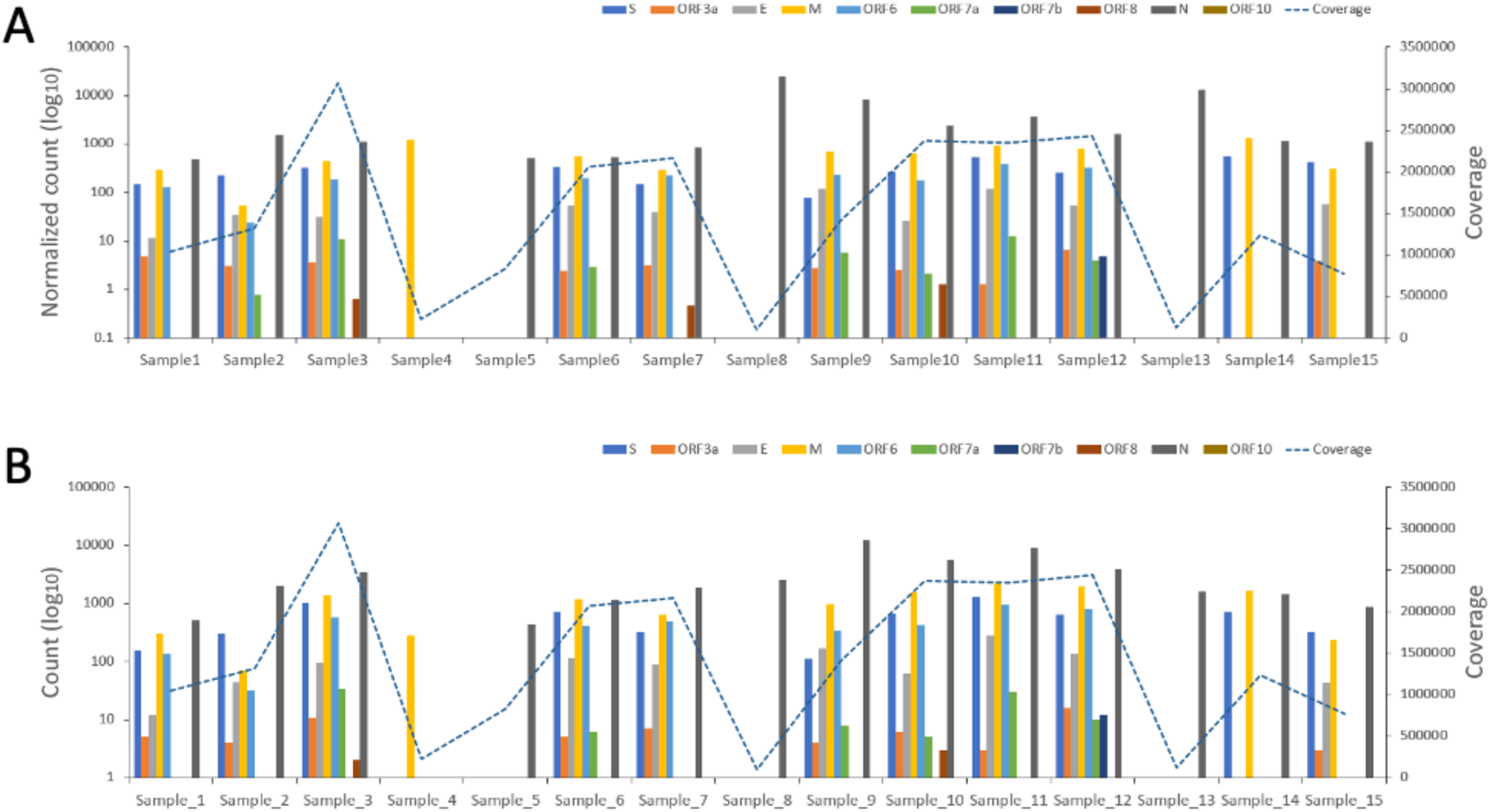
Plots of normalised peak counts (A) and peak counts (B) of leader-TRS gene junctions of reads with at least one primer sequences at either end derived from sequence data from 15 human patients. These were sequenced with the ARTIC pipeline via Illumina. The data correspond to Supplementary Table 7.

**Figure 5.**
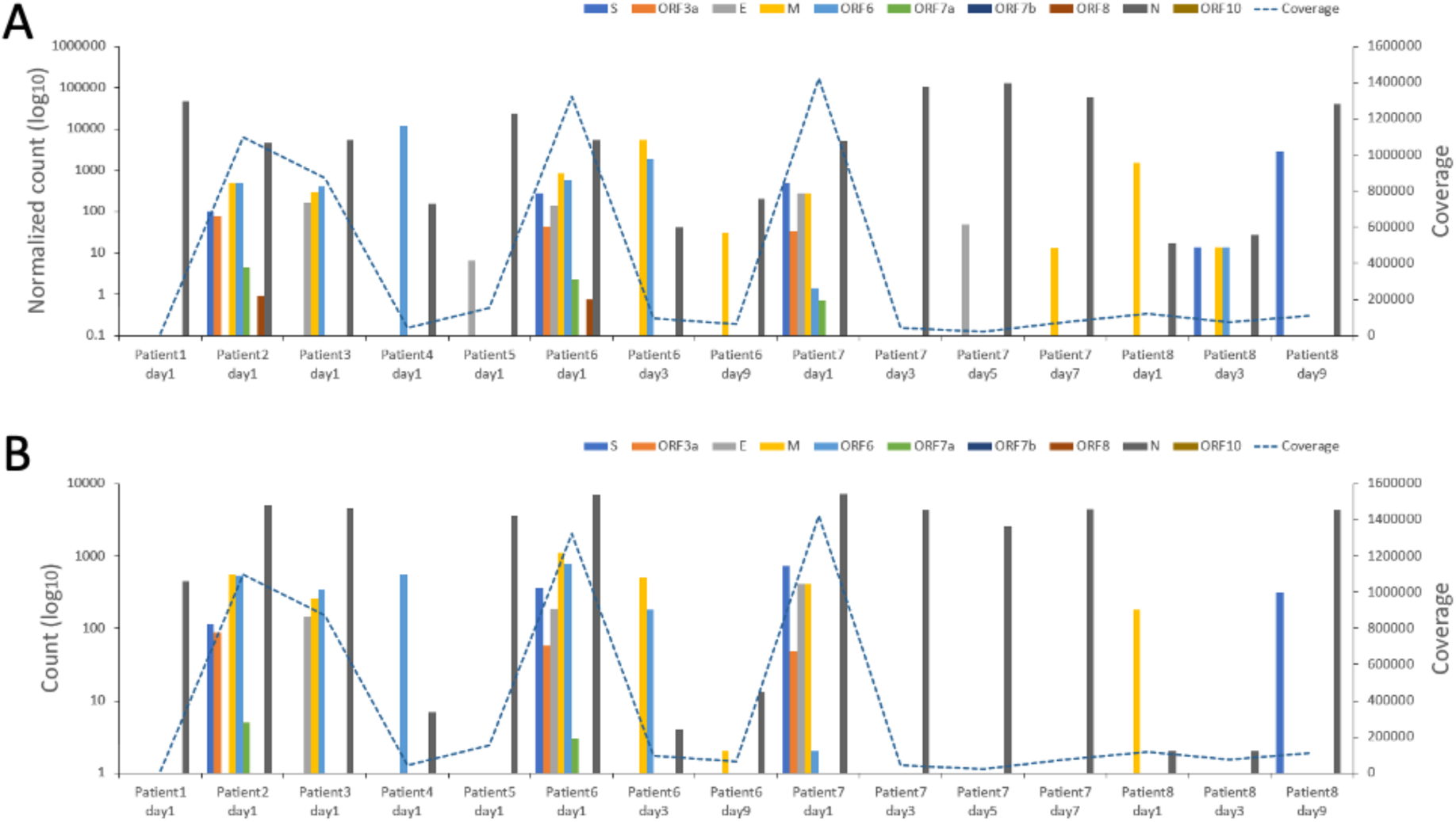
Plots of normalised peak counts (A) and peak counts (B) of leader-TRS gene junctions of reads with at least one primer sequence at either end derived from sequence data from 15 human patients. Some of these samples are longitudinal as indicated by the patient number and day post admission the sample was taken. These were sequenced with the ARTIC pipeline via Nanopore. The data are correspond to Supplementary Table 10.

### Commonality of novel leader-TRS gene junctions

The sequencing data spanning cell culture infection, animal models and clinical samples from humans indicated the presence of novel leader-TRS gene junctions. Their detection generally increased with depth of coverage. Coronavirus replication and transcription is promiscuous, and recombination is a natural result of this, resulting in insertions, deletions and potential gene rearrangements. Many of these novel leader-TRS junctions were centred around the known gene open reading frame but out of the search interval. These type of leader-TRS gene junctions could be only found with spike, membrane, ORF6, ORF7b and nucleocapsid orfs, in which the membrane orf was the most common (Figure 6A). In order to define what might be genuine novel leader-TRS gene junctions, these were compared across the data in all Illumina ARTIC data (Figure 6B, Supplementary Table 11). This identified 5 novel leader-TRS junctions that were common to all the data, the majority of these being focused on the membrane orf.

**Figure 6.**
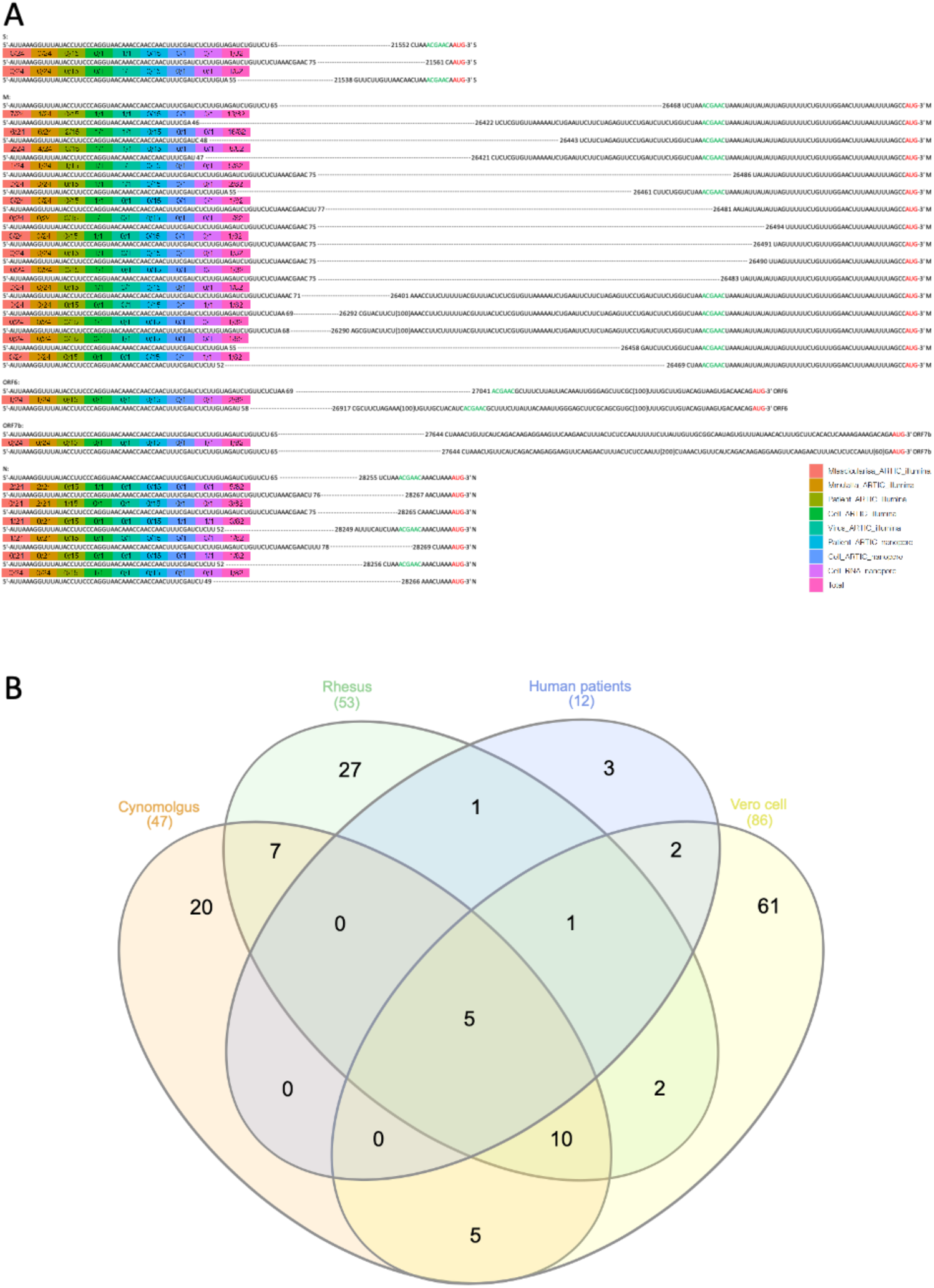
(A). Diagram of novel leader-TRS junctions centred around the known gene orf but out of the search interval in the analysis of cell culture, non-human primate and human sequencing data. Many novel junctions map to the leader-TRS membrane gene junctions. (B). Venn diagram showing the overlap of novel leader-TRS gene junctions among cynomolgus and rhesus macaques, human patients, and infected Vero cells all sequenced with the ATRIC Illumina method (Supplementary Table 11).

## Discussion

Coronavirus sgmRNAs are only synthesised during infection of cells and therefore their presence in sequence data can be indicative of active viral RNA synthesis. The abundance of the sgmRNAs in infected cells should follow a general pattern where the sgmRNA encoding the nucleoprotein is the most abundant. Identification and quantification of the unique leader-TRS gene junctions for each sgmRNA can be used as a proxy for their abundance.

LeTRS was developed to interrogate sequencing datasets to identify the leader-gene junctions present at the 5’ end of the sgmRNAs. LeTRS was first evaluated and validated on cell culture data from published datasets^2, 16^ and from a cell culture experiment as part of this study and then used in an analysis of nasopharyngeal samples from non-human primate and human clinical samples. The results showed that the positions of the leader-TRS junction sites with peak read counts were same as the given reference positions. The exception was at leader-gene junction for orf7b in the Nanopore sequencing. The normalized count results confirmed the reads spanning the junctions showed that the leader-TRS nucleoprotein gene junction was the most abundant, and orf7b and orf10 were the most infrequent in line with other data^2, 20^. Several low abundant leader-TRS junctions were identified in all of the datasets with the implication these were either from potential lower abundant novel sgmRNAs, or represented known sgmRNAs, but with different leader-TRS junctions. Likewise, at low frequency these could represent an aberrant viral transcription process or artefacts of the different sequencing processes – although this latter possibility is less likely through the published direct RNA sequencing approach^2^ (Figure 2B). Traditionally, such sgmRNAs have been first identified in coronaviruses by either northern blot and/or metabolic labelling^8^. Several other groups have identified novel leader-TRS gene junctions and potential subgenomic mRNAs for other coronaviruses, including avian infectious bronchitis virus^21^. The best way of validating potential novel sgmRNAs would be through matching proteomic data to confirm genuine open reading frames^1^. Analysis of several published sequencing datasets identified novel viral RNA molecules that the authors suggested were sgmRNAs containing only the 5’ region of orf1a^22^. Such species are likely to be defective RNAs, that act as templates for replication. Interestingly, they hypothesize that at later time points post-infection in cell culture potential novel sgRNAs are generated non-specifically^22^, which potentially ties in with a disconnect of leader-TRS gene junctions observed in our study both *in vivo* from the nasopharyngeal samples from latter time points in the NHP models and in humans and from the published data from SARS-CoV-2 infections in cell culture gathered at later time points compared to earlier time points^2, 16^.

Advanced filtering can improve the confidence of the identified leader-TRS junction in the sequencing reads. Amplicon sequencing provided a unique opportunity to filter the sequencing reads. The reads spanning the junctions with the correct forward primer, reverse primer or both primer sequences at the ends of reads proved the known/novel sgmRNA existing in tested Illumina and Nanopore ARTIC v3 primers amplicon sequencing data (Tables 1 and 3). For Illumina sequencing, the same junction on paired reads with at least a primer provided extra evidence for leader-TRS identification. Some reads were identified that did not have primer sequences and these were likely to be miss-mapped, from template sgmRNA or low-quality sequence. These were present at very low abundance compared to authentic mapped reads (Tables 1, 3 and 5). No reads with polyA were detected in the Nanopore amplicon sequencing data, this was likely because the limited PCR extension time restricted the primers to reach the 5’end of subgenomic mRNA (Table 1 and Supplementary Table5). The Nanopore direct RNA sequencing had the potential to generate full length mRNA sequences. The polyA sequences and leader-TRS junctions in the reads can be good signals to prove the full length sgmRNA in the test data (Tables 3 and 4). Because the fortuitous sequencing of some host mRNA may lead false positive result, LeTRS was tested against sequence data from uninfected controls cells^2^. No positive reads were found in this control sample (Tables 5 and 6), suggesting the LeTRS could effectively screen out or not recognise any false positives. Crucially, LeTRS used less CPU runtime and provided more detailed information than other tools to investigate this^17^, and therefore is suited for the high throughput analysis of large amounts of diverse sequencing data.

**Table 4.** The LeTRS output table for novel sgmRNA in the tested Nanopore direct RNA sequencing data. “leader_end” and “TRS_start” refer to the position of the end of leader and the position of the start of TRS identified in the reads >10.

| subgenome | leader_end | TRS_start | nb_count | normalized_count |
| --- | --- | --- | --- | --- |
| 1 | 52 | 26469 | 40(36,13) | 69.49(62.54,22.59) |
| 2 | 52 | 28249 | 99(90,36) | 172.00(156.36,62.54) |
| 3 | 52 | 28256 | 138(127,50) | 239.75(220.64,86.87) |
| 4 | 60 | 21045 | 16(16,6) | 27.80(27.80,10.42) |
| 5 | 65 | 22273 | 17(16,5) | 29.53(27.80,8.69) |
| 6 | 65 | 22488 | 12(11,5) | 20.85(19.11,8.69) |
| 7 | 65 | 24777 | 24(23,11) | 41.70(39.96,19.11) |
| 8 | 65 | 27479 | 51(48,18) | 88.60(83.39,31.27) |
| 9 | 69 | 21053 | 17(14,6) | 29.53(24.32,10.42) |
| 10 | 69 | 26292 | 14(14,4) | 24.32(24.32,6.95) |
| 11 | 69 | 28284 | 14(14,7) | 24.32(24.32,12.16) |
| 12 | 69 | 29112 | 21(21,6) | 36.48(36.48,10.42) |
| 13 | 69 | 29152 | 11(11,4) | 19.11(19.11,6.95) |

The numbers in the bracket are (reads with > 1 poly A, reads with > 5 poly A).
Normalized count=(Read count/Total number of read mapped on reference genome)\*1000000.
Total number of read mapped on reference genome is 575596, excluding mapped reads on reverse strand, not primary alignment and supplementary alignment.

In terms of clinical samples (typically nasopharyngeal swabs), the presence of sgmRNAs will be due to the presence infected cells. In general, this has been seen as indicative of active viral RNA synthesis at the time of sampling^5, 23, 24^, although these have also been postulated to be present through resistant structures after infection has finished^25^. Analysis of inoculum indicated that leader-TRS gene junctions could be identified (Supplementary Figure 3) but that these were not in the same ratio as found in cells infected in culture (e.g., Figure 2B and 2C). Thus, if the abundance of leader-TRS gene junctions follows an expected pattern of the nucleoprotein gene leader-TRS gene junction being the most abundant followed by a general gradient in sequence data from nasopharyngeal samples, then this may be indicative of an active infection – and the presence of infected cells in a sample.

In the absence of a human challenge model, NHP models that closely resemble COVID-19 disease in humans can be used to study SARS-CoV-2 infection, from a very defined initial exposure. RNA was sequenced from longitudinal nasopharyngeal samples from two NHP models, rhesus and cynomolgus macaques^18^. LeTRS used to identify the abundance of the leader-TRS gene junctions in this data. The analysis indicated a phasic pattern of sgmRNA synthesis with a large drop off after Day 8/9 post-infection in both NHP models. This phasic pattern may be explained by an initial synchronous infection of respiratory epithelial cells and then these cells dying. Released virus then goes on to infect new epithelial cells with virus infection increasing exponentially in waves but becoming asynchronous. The decline in sgmRNA from Day 8/9 overlaps with IgG seroconversion and humoral immunity in both species^18^ and follows similar kinetics to serology profiles measured in patients with COVID-19.

The identification of sgmRNAs in nasopharyngeal samples and their kinetics has implications for nucleic acid-based diagnostics (many of which have three targets, one in the orf1a/b region and two which are shared between the genome and sgmRNAs – the nucleoprotein and the spike genes). The phasic nature of leader-TRS gene junctions in the longitudinal samples, and by implication sgmRNAs, and overt abundance of the leader-TRS nucleoprotein gene junction found in many of the human samples, suggests that it may not be possible to precisely identify where in infection an individual is based on the abundance of sgmRNAs. Likewise, assuming equivalency between the targets, if the nucleoprotein target is found to be more abundant than the spike target than the genomic target, then this would suggest infected cells are present in the sample. Decreases in Ct values associated with emerging variants could equally be explained by sloughed cells being present in a nasopharyngeal sample as well as by increases in the amount of virions/viral load. Therefore, we would caution that a decrease in Ct associated with RT-qPCR based assays may not just be reflective of higher viral loads but also may be indicative of more infected cells being present. These possibilities may be resolved by considering the relative ratios of sgmRNAs identified.

## METHODS

### Data input

LeTRS was designed to analyse FastQ files derived from Illumina paired-end or Nanopore sequencing data derived from a SARS-CoV-2 amplicon protocol, or standard Nanopore SARS-CoV-2 direct RNA sequencing data (Figure 1). The Illumina/Nanopore FastQ sequencing data were cleaned to remove adapters and low-quality reads before input. Sequencing data derived from other sequencing modes or platforms can also be analysed by LeTRS via input of a BAM file produced by a custom splicing alignment method with a SARS-CoV-2 genome (NC_045512.2) as a reference (Figure 1). This can also be rapidly adapted for other coronaviruses.

### Library preparations and sequencing

We sequenced the 15 samples from human patients with nanopore. Total RNA was isolated using a QIAamp Viral RNA Mini Kit (Qiagen, Manchester, UK) by spin-column procedure according to the manufacturer’s instructions. Clinical samples were extracted with Trizol LS as described^4^. All RNA samples were treated with Turbo DNase (Invitrogen). SuperScript IV (Invitrogen) was used to generate single-strand cDNA using random primer mix (NEB, Hitchin, UK). ARTIC V3 PCR amplicons from the single-strand cDNA were generated following the Nanopore Protocol of PCR tiling of SARS-CoV-2 virus (Version: PTC_9096_v109_revL_06Feb2020). Amplicons generated by ARTIC PCR were purified and normalised to 200 fmol before DNA end preparation and barcode and adapter ligation. Library was loaded onto a FLO-MIN106 flow cell and sequencing reads were called with Guppy using the high-accuracy calling parameters.

The NHP samples and their inoculum, and our laboratory experiments conducted in cells were sequenced with Illumina. The amplicons products for Illumina sequencing were prepared as per the Nanopore sequencing above and then used in Illumina NEBNext Ultra II DNA Library preparation. Following 4 cycles of amplification the library was purified using Ampure XP beads and quantified using Qubit and the size distribution assessed using the Fragment analyzer. Finally, the ARTIC library was sequenced on the Illumina® NovaSeq 6000 platform (Illumina®, San Diego, USA) following the standard workflow. The generated raw FastQ files (2 x 250 bp) were trimmed to remove Illumina adapter sequences using Cutadapt v1.2.1 ^26^. The option “−O 3” was set, so the that 3’ end of any reads which matched the adapter sequence with greater than 3 bp was trimmed off. The reads were further trimmed to remove low quality bases, using Sickle v1.200 ^27^ with a minimum window quality score of 20. After trimming, reads shorter than 10 bp were removed.

The LeTRS was also tested with a combined Nanopore cDNA ARTIC v3 amplicon dataset of 7 published viral cell culture samples (barcode01-barcode07) ^16^, and a dataset from a published direct RNA Nanopore sequencing analysis Vero cells infected with SARS-CoV-2 or an uninfected negative control ^2^.

### Sequencing data alignment and basic filtering

LeTRS controlled Hisat2 v2.1.0 ^28^ to map the paired-end Illumina reads against the SARS-CoV-2 reference genome (NC_045512.2) with the default setting, and Minimap2 v2.1 ^29^ to align the Nanopore cDNA reads and direct RNA-seq reads on the viral genome using Minimap2 with “–ax splice” and “-ax splice -uf -k14” parameters, respectively. LeTRS provided 10 known leader-TRS junctions to improve alignment accuracy by using “--known-splicesite-infile” function in Hisat2 and “--junc-bed” function in Minimap2, but this application could be optionally switched off by users. In order to remove low mapping quality and mis-mapped reads before searching the leader-TRS junction sites, LeTRS used Samtools v1.9 ^30^ to have basic filtering for the reads in the output Sam/Bam files according to their alignment states as shown (Table 9 - basic filtering).

**Table 9.**
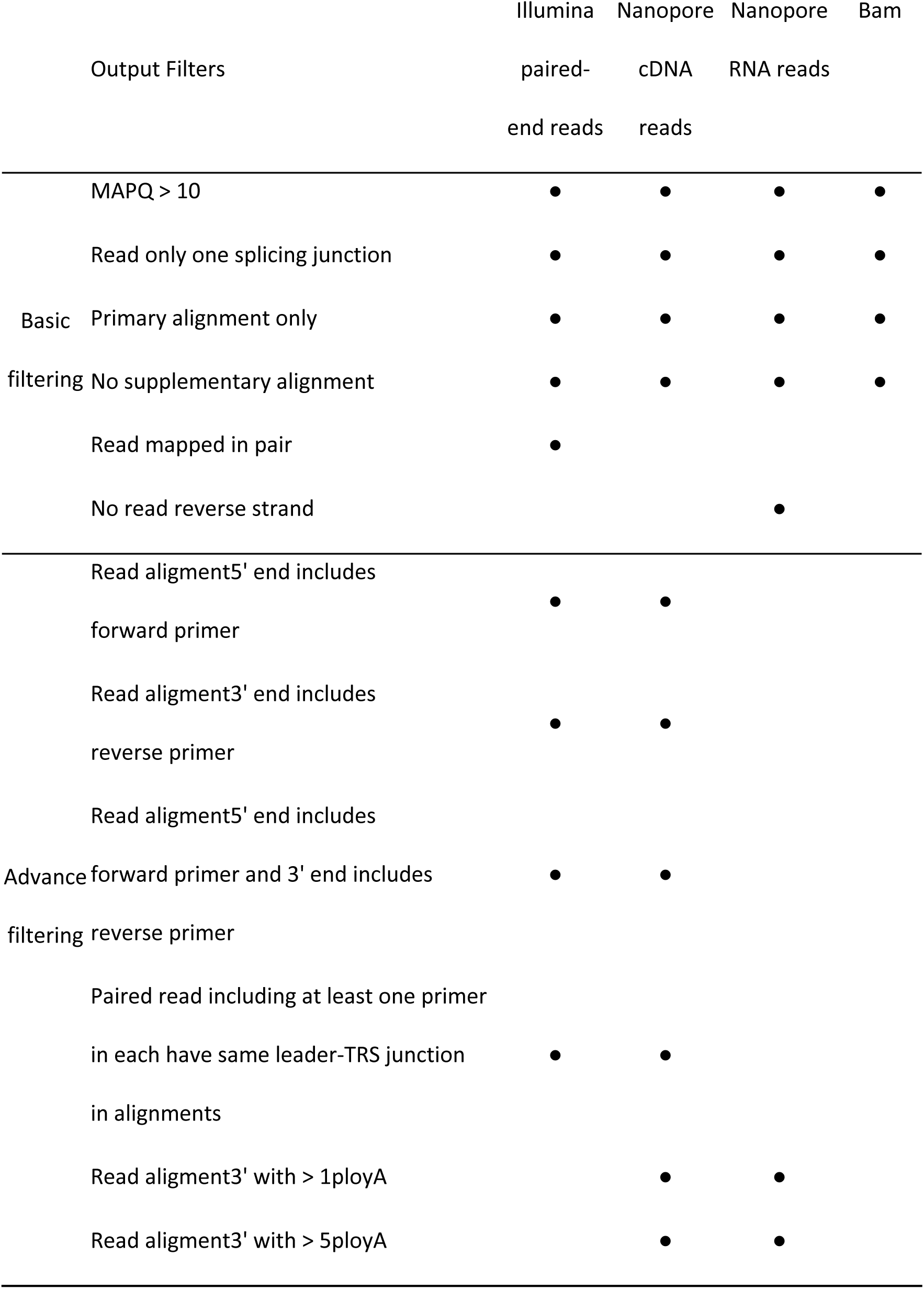
The criteria of basic and advanced filtering for four different types of input data for LeTRS.

### Search leader-TRS

After the mapping and basic filtering step, LeTRS searched aligned reads spanning the leader-TRS junctions in the SARS-CoV-2 reference genome (Supplementary Figure 1). For the known leader-TRS junctions, LeTRS searched the reads including the leader-TRS junctions within a given interval around the known leader and TRS junctions sites. The leader break site interval is ±10 nts, and the TRS breaking sites interval is -20 nts to the 1 nt before the first known AUG in the default setting (the intervals can be changed to custom values to investigate heterogeneity). LeTRS then reported a peak count that was the number of reads carrying the most common leader-TRS junctions within the given leader and TRS breaking sites intervals, and a cluster count that was the number of all reads carrying leader-TRS junctions within the given leader and TRS breaking sites intervals (Tables 1-6). LeTRS also searched the junctions out of the given intervals (the genomic position of leader breaking site < 80) and reported the number of reads (>10 by default) with novel leader-TRS junctions. These number of read counts were also reported by number of reads in 1000000 as normalization. The read including the known and novel leader-TRS junctions could be optionally outputted in FastA format. Based on identified known and novel leader-TRS junctions, LeTRS could report 20 nucleotides towards the 3’ end of the leader sequence, the TRS and translated the first orf of sgmRNAs sequence, and find the conserved ACGAAC sequences in the TRS (Table S1-S6).

### Advance filtering

Based on the alignment possibilities illustrated in Figure 2 and discussed, LeTRS further filters the identified reads with known and novel leader-TRS junctions. This step is named as advance filtering and can only applied when the input data is from Illumina paired end reads, Nanopore cDNA reads or Nanopore RNA reads (Table 9). If a BAM file is used as input data, the advanced filtering step would be automatically skipped (Table 9). The number of reads including the known and novel leader-TRS junctions, and the number of reads filtered with corresponding advance filtering criteria were outputted into two tables in tab format (Tables 1-6).

### Leader-TRS junction plotting

LeTRS-plot was developed as an automatic plotting tool that interfaces with the R package ggplot2 v3.3.3 to view the leader-TRS junctions in the tables generated by LeTRS (Figure 3-5). The plot shows peak count, filtered peak count, normalized peak count and normalized filtered peak count for known leader-TRS junctions, and novel junction counts, filtered novel junction count, normalized novel junction count and filtered normalized novel junction for novel leader-TRS junctions.

## Ethics approval and consent to participate

All experimental work on NHPs was conducted under the authority of a UK Home Office approved project license (PDC57C033) that had been subject to local ethical review at PHE Porton Down by the Animal Welfare and Ethical Review Body (AWERB) and approved as required by the Home Office Animals (Scientific Procedures) Act 1986 and the full ethics and NHP model are described.

## Consent for publication

Not applicable

## Availability of data and materials

LeTRS is available at https://github.com/xiaofengdong83/LeTRS.

Illumina and nanopore test data sets are available under NCBI PRJNA699398.

## Authors’ contributions

X.D. developed the LeTRS software and performed the informatics analysis. X.D., A.D. and J.A.H. analysed the data. J.S., J.T. and M.W.C. co-ordinated the NHP work and sample processing. R.P.-R., J.P.S., H.G., T.P. and N.R. were involved in sequencing and informatics analysis of the NHP samples with D.A.M. A.D. oversaw sequencing of the human clinical samples with E.V. and C.N for the COG-UK data. R.P.-R. and J.A.H. oversaw sequencing of samples under the auspices of ISARIC-4C managed by J.K.B, L.T., M.G.S. and P.J.M.O. J.A.H. and M.W.C. initiated and managed the study and wrote the manuscript with X.D., R.P.-R., A.D. with other authors involved in editing the final version.

## Acknowledgments

We would like to thank all members of the Hiscox Laboratory and the Centre for Genome Research for supporting SARS-CoV-2/COVID-19 sequencing research.

## Consortia

**The members of the COG-UK consortia are:** Thomas R. Connor, Nicholas J. Loman, Samuel C. Robson, Tanya Golubchik, M. Estee Torok, William L. Hamilton, David Bonsall, Ali R. Awan, Sally Corden, Ian Goodfellow, Darren L. Smith, Martin D. Curran, Surendra Parmar, James G. Shepherd, Matthew D. Parker, Catherine Moore, Derek J. Fairley, Matthew W. Loose, Joanne Watkins, Matthew Bull, Sam Nicholls, David M. Aanensen, Sharon Glaysher, Matthew Bashton, Nicole Pacchiarini, Anthony P. Underwood, Thushan I. de Silva, Dennis Wang, Monique Andersson, Anoop J. Chauhan, Mariateresa de Cesare, Catherine Ludden, Tabitha W. Mahungu, Rebecca Dewar, Martin P. McHugh, Natasha G. Jesudason, Kathy K. Li, Rajiv N. Shah, Yusri Taha, Kate E. Templeton, Simon Cottrell, Justin O’Grady, Andrew Rambaut, Colin P. Smith, Matthew T.G. Holden, Emma C. Thomson, Samuel Moses, Meera Chand, Chrystala Constantinidou, Alistair C. Darby, Julian A. Hiscox, Steve Paterson, Meera Unnikrishnan, Andrew J. Page, Erik M. Volz, Charlotte J. Houldcroft, Aminu S. Jahun, James P. McKenna, Luke W. Meredith, Andrew Nelson, Sarojini Pandey, Gregory R. Young, Anna Price, Sara Rey, Sunando Roy, Ben Temperton, Matthew Wyles, Stefan Rooke, Sharif Shaaban, Helen Adams, Yann Bourgeois, Katie F. Loveson, Áine O’Toole, Richard Stark, Ewan M. Harrison, David Heyburn, Sharon J. Peacock, David Buck, Michaela John, Dorota Jamrozy, Joshua Quick, Rahul Batra, Katherine L. Bellis, Beth Blane, Sophia T. Girgis, Angie Green, Anita Justice, Mark Kristiansen, Rachel J. Williams, Radoslaw Poplawski, Garry P. Scarlett, John A. Todd, Christophe Fraser, Judith Breuer, Sergi Castellano, Stephen L. Michell, Dimitris Gramatopoulos, Jonathan Edgeworth, Gemma L. Kay, Ana da Silva Filipe, Aaron R. Jeffries, Sascha Ott, Oliver Pybus, David L. Robertson, David A. Simpson, Chris Williams, Cressida Auckland, John Boyes, Samir Dervisevic, Sian Ellard, Sonia Goncalves, Emma J. Meader, Peter Muir, Husam Osman, Reenesh Prakash, Venkat Sivaprakasam, Ian B. Vipond, Jane A.H. Masoli, Nabil-Fareed Alikhan, Matthew Carlile, Noel Craine, Sam T. Haldenby, Nadine Holmes, Ronan A. Lyons, Christopher Moore, Malorie Perry, Ben Warne, Thomas Williams, Lisa Berry, Andrew Bosworth, Julianne Rose Brown, Sharon Campbell, Anna Casey, Gemma Clark, Jennifer Collins, Alison Cox, Thomas Davis, Gary Eltringham, Cariad Evans, Clive Graham, Fenella Halstead, Kathryn Ann Harris, Christopher Holmes, Stephanie Hutchings, Miren Iturriza-Gomara, Kate Johnson, Katie Jones, Alexander J. Keeley, Bridget A. Knight, Cherian Koshy, Steven Liggett, Hannah Lowe, Anita O. Lucaci, Jessica Lynch, Patrick C McClure, Nathan Moore, Matilde Mori, David G. Partridge, Pinglawathee Madona, Hannah M. Pymont, Paul Anthony Randell, Mohammad Raza, Felicity Ryan, Robert Shaw, Tim J. Sloan, Emma Swindells, Alexander Adams, Hibo Asad, Alec Birchley, Tony Thomas Brooks, Giselda Bucca, Ethan Butcher, Sarah L. Caddy, Laura G. Caller, Yasmin Chaudhry, Jason Coombes, Michelle Cronin, Patricia L. Dyal, Johnathan M. Evans, Laia Fina, Bree Gatica-Wilcox, Iliana Georgana, Lauren Gilbert, Lee Graham, Danielle C. Groves, Grant Hall, Ember Hilvers, Myra Hosmillo, Hannah Jones, Sophie Jones, Fahad A. Khokhar, Sara Kumziene-Summerhayes, George MacIntyre-Cockett, Rocio T. Martinez Nunez, Caoimhe McKerr, Claire McMurray, Richard Myers, Yasmin Nicole Panchbhaya, Malte L. Pinckert, Amy Plimmer, Joanne Stockton, Sarah Taylor, Alicia Thornton, Amy Trebes, Alexander J. Trotter, Helena Jane Tutill, Charlotte A. Williams, Anna Yakovleva, Wen C. Yew, Mohammad T. Alam, Laura Baxter, Olivia Boyd, Fabricia F. Nascimento, Timothy M. Freeman, Lily Geidelberg, Joseph Hughes, David Jorgensen, Benjamin B. Lindsey, Richard J. Orton, Manon Ragonnet-Cronin, Joel Southgate, Sreenu Vattipally, Igor Starinskij, Joshua B. Singer, Khalil Abudahab, Leonardo de Oliveira Martins, Thanh Le-Viet, Mirko Menegazzo, Ben E.W. Taylor, Corin A. Yeats, Sophie Palmer, Carol M. Churcher, Alisha Davies, Elen De Lacy, Fatima Downing, Sue Edward, Nikki Smith, Frances Bolt, Alex Alderton, Matt Berriman, Ian G. Charles, Nicholas Cortes, Tanya Curran, John Danesh, Sahar Eldirdiri, Ngozi Elumogo, Andrew Hattersley, Alison Holmes, Robin Howe, Rachel Jones, Anita Kenyon, Robert A. Kingsley, Dominic Kwiatkowski, Cordelia Langford, Jenifer Mason, Alison E. Mather, Lizzie Meadows, Sian Morgan, James Price, Trevor I. Robinson, Giri Shankar, John Wain, Mark A. Webber, Declan T. Bradley, Michael R. Chapman, Derrick Crooke, David Eyre, Martyn Guest, Huw Gulliver, Sarah Hoosdally, Christine Kitchen, Ian Merrick, Siddharth Mookerjee, Robert Munn, Timothy Peto, Will Potter, Dheeraj K Sethi, Wendy Smith, Luke B. Snell, Rachael Stanley, Claire Stuart, Elizabeth Wastenge, Erwan Acheson, Safiah Afifi, Elias Allara, Roberto Amato, Adrienn Angyal, Elihu Aranday-Cortes, Cristina Ariani, Jordan Ashworth, Stephen Attwood, Alp Aydin, David J. Baker, Carlos E. Balcazar, Angela Beckett, Robert Beer, Gilberto Betancor, Emma Betteridge, David Bibby, Daniel Bradshaw, Catherine Bresner, Hannah E. Bridgewater, Alice Broos, Rebecca Brown, Paul E. Brown, Kirstyn Brunker, Stephen N. Carmichael, Jeffrey K.J. Cheng, Rachel Colquhoun, Gavin Dabrera, Johnny Debebe, Eleanor Drury, Louis du Plessis, Richard Eccles, Nicholas Ellaby, Audrey Farbos, Ben Farr, Jacqueline Findlay, Chloe L. Fisher, Leysa Marie Forrest, Sarah Francois, Lucy R. Frost, William Fuller, Eileen Gallagher, Michael D. Gallagher, Matthew Gemmell, Rachel A.J. Gilroy, Scott Goodwin, Luke R. Green, Richard Gregory, Natalie Groves, James W. Harrison, Hassan Hartman, Andrew R. Hesketh, Verity Hill, Jonathan Hubb, Margaret Hughes, David K. Jackson, Ben Jackson, Keith James, Natasha Johnson, Ian Johnston, Jon-Paul Keatley, Moritz Kraemer, Angie Lackenby, Mara Lawniczak, David Lee, Rich Livett, Stephanie Lo, Daniel Mair, Joshua Maksimovic, Nikos Manesis, Robin Manley, Carmen Manso, Angela Marchbank, Inigo Martincorena, Tamyo Mbisa, Kathryn McCluggage, J.T. McCrone, Shahjahan Miah, Michelle L. Michelsen, Mari Morgan, Gaia Nebbia, Charlotte Nelson, Jenna Nichols, Paola Niola, Kyriaki Nomikou, Steve Palmer, Naomi Park, Yasmin A. Parr, Paul J. Parsons, Vineet Patel, Minal Patel, Clare Pearson, Steven Platt, Christoph Puethe, Mike Quail, Jayna Raghwani, Lucille Rainbow, Shavanthi Rajatileka, Mary Ramsay, Paola C. Resende Silva, Steven Rudder, Chris Ruis, Christine M. Sambles, Fei Sang, Ulf Schaefer, Emily Scher, Carol Scott, Lesley Shirley, Adrian W. Signell, John Sillitoe, Christen Smith, Katherine L. Smollett, Karla Spellman, Thomas D. Stanton, David J. Studholme, Grace Taylor-Joyce, Ana P. Tedim, Thomas Thompson, Nicholas M. Thomson, Scott Thurston, Lily Tong, Gerry Tonkin-Hill, Rachel M. Tucker, Edith E. Vamos, Tetyana Vasylyeva, Joanna Warwick-Dugdale, Danni Weldon, Mark Whitehead, David Williams, Kathleen A. Williamson, Harry D. Wilson, Trudy Workman, Muhammad Yasir, Xiaoyu Yu, Alex Zarebski, Evelien M. Adriaenssens, Shazaad S.Y. Ahmad, Adela Alcolea-Medina, John Allan, Patawee Asamaphan, Laura Atkinson, Paul Baker, Jonathan Ball, Edward Barton, Mathew A. Beale, Charlotte Beaver, Andrew Beggs, Andrew Bell, Duncan J Berger, Louise Berry, Claire M. Bewshea, Kelly Bicknell, Paul Bird, Chloe Bishop, Tim Boswell, Cassie Breen, Sarah K. Buddenborg, Shirelle Burton-Fanning, Vicki Chalker, Joseph G. Chappell, Themoula Charalampous, Claire Cormie, Nick Cortes, Lindsay J. Coupland, Angela Cowell, Rose K. Davidson, Joana Dias, Maria Diaz, Thomas Dibling, Matthew J. Dorman, Nichola Duckworth, Scott Elliott, Sarah Essex, Karlie Fallon, Theresa Feltwell, Vicki M Fleming, Sally Forrest, Luke Foulser, Maria V. Garcia-Casado, Artemis Gavriil, Ryan P. George, Laura Gifford, Harmeet K. Gill, Jane Greenaway, Luke Griffith, Ana Victoria Gutierrez, Antony D. Hale, Tanzina Haque, Katherine L. Harper, Ian Harrison, Judith Heaney, Thomas Helmer, Ellen E. Higginson, Richard Hopes, Hannah C. Howson-Wells, Adam D. Hunter, Robert Impey, Dianne Irish-Tavares, David A. Jackson, Kathryn A. Jackson, Amelia Joseph, Leanne Kane, Sally Kay, Leanne M. Kermack, Manjinder Khakh, Stephen P. Kidd, Anastasia Kolyva, Jack C.D. Lee, Laura Letchford, Nick Levene, Lisa J. Levett, Michelle M. Lister, Allyson Lloyd, Joshua Loh, Louissa R. Macfarlane-Smith, Nicholas W. Machin, Mailis Maes, Samantha McGuigan, Liz McMinn, Lamia Mestek-Boukhibar, Zoltan Molnar, Lynn Monaghan, Catrin Moore, Plamena Naydenova, Alexandra S. Neaverson, Rachel Nelson, Marc O. Niebel, Elaine O’Toole, Debra Padgett, Gaurang Patel, Brendan A.I. Payne, Liam Prestwood, Veena Raviprakash, Nicola Reynolds, Alex Richter, Esther Robinson, Hazel A. Rogers, Aileen Rowan, Garren Scott, Divya Shah, Nicola Sheriff, Graciela Sluga, Emily Souster, Michael Spencer-Chapman, Sushmita Sridhar, Tracey Swingler, Julian Tang, Graham P. Taylor, Theocharis Tsoleridis, Lance Turtle, Sarah Walsh, Michelle Wantoch, Joanne Watts, Sheila Waugh, Sam Weeks, Rebecca Williams, Iona Willingham, Emma L. Wise, Victoria Wright, Sarah Wyllie, Jamie Young, Amy Gaskin, Will Rowe, Igor Siveroni, and Robert Johnson.

**The members of the ISARIC4C consortia are:** Consortium Lead Investigator: J. Kenneth Baillie; Chief Investigator: Malcolm G. Semple; Co-Lead Investigator: Peter J.M. Openshaw; ISARIC Clinical Coordinator: Gail Carson; Co-Investigators: Beatrice Alex, Benjamin Bach, Wendy S. Barclay, Debby Bogaert, Meera Chand, Graham S. Cooke, Annemarie B. Docherty, Jake Dunning, Ana da Silva Filipe, Tom Fletcher, Christopher A. Green, Ewen M. Harrison, Julian A. Hiscox, Antonia Ying Wai Ho, Peter W. Horby, Samreen Ijaz, Saye Khoo, Paul Klenerman, Andrew Law, Wei Shen Lim, Alexander J. Mentzer, Laura Merson, Alison M. Meynert, Mahdad Noursadeghi, Shona C. Moore, Massimo Palmarini, William A. Paxton, Georgios Pollakis, Nicholas Price, Andrew Rambaut, David L. Robertson, Clark D. Russell, Vanessa Sancho-Shimizu, Janet T. Scott, Thushan de Silva, Louise Sigfrid, Tom Solomon, Shiranee Sriskandan, David Stuart, Charlotte Summers, Richard S. Tedder, Emma C. Thomson, A.A. Roger Thompson, Ryan S. Thwaites, Lance C.W. Turtle, and Maria Zambon; Project Managers: Hayley Hardwick, Chloe Donohue, Ruth Lyons, Fiona Griffiths, and Wilna Oosthuyzen; Data Analysts: Lisa Norman, Riinu Pius, Tom M. Drake, Cameron J. Fairfield, Stephen Knight, Kenneth A. Mclean, Derek Murphy, and Catherine A. Shaw; Data and Information System Managers: Jo Dalton, James Lee, Daniel Plotkin, Michelle Girvan, Egle Saviciute, Stephanie Roberts, Janet Harrison, Laura Marsh, Marie Connor, Sophie Halpin, Clare Jackson, and Carrol Gamble; Data Integration and Presentation: Gary Leeming, Andrew Law, Murray Wham, Sara Clohisey, Ross Hendry, and James Scott-Brown; Material Management: William Greenhalf, Victoria Shaw, and Sarah McDonald; Patient Engagement: Seán Keating; Outbreak Laboratory Staff and Volunteers: Katie A. Ahmed, Jane A. Armstrong, Milton Ashworth, Innocent G. Asiimwe, Siddharth Bakshi, Samantha L. Barlow, Laura Booth, Benjamin Brennan, Katie Bullock, Benjamin W.A. Catterall, Jordan J. Clark, Emily A. Clarke, Sarah Cole, Louise Cooper, Helen Cox, Christopher Davis, Oslem Dincarslan, Chris Dunn, Philip Dyer, Angela Elliott, Anthony Evans, Lorna Finch, Lewis W.S. Fisher, Terry Foster, Isabel Garcia-Dorival, Willliam Greenhalf, Philip Gunning, Catherine Hartley, Antonia Ho, Rebecca L. Jensen, Christopher B. Jones, Trevor R. Jones, Shadia Khandaker, Katharine King, Robyn T. Kiy, Chrysa Koukorava, Annette Lake, Suzannah Lant, Diane Latawiec, L. Lavelle-Langham, Daniella Lefteri, Lauren Lett, Lucia A. Livoti, Maria Mancini, Sarah McDonald, Laurence McEvoy, John McLauchlan, Soeren Metelmann, Nahida S. Miah, Joanna Middleton, Joyce Mitchell, Shona C. Moore, Ellen G. Murphy, Rebekah Penrice-Randal, Jack Pilgrim, Tessa Prince, Will Reynolds, P. Matthew Ridley, Debby Sales, Victoria E. Shaw, Rebecca K. Shears, Benjamin Small, Krishanthi S. Subramaniam, Agnieska Szemiel, Aislynn Taggart, Jolanta Tanianis-Hughes, Jordan Thomas, Erwan Trochu, Libby van Tonder, Eve Wilcock, and J. Eunice Zhang; Local Principal Investigators: Kayode Adeniji, Daniel Agranoff, Ken Agwuh, Dhiraj Ail, Ana Alegria, Brian Angus, Abdul Ashish, Dougal Atkinson, Shahedal Bari, Gavin Barlow, Stella Barnass, Nicholas Barrett, Christopher Bassford, David Baxter, Michael Beadsworth, Jolanta Bernatoniene, John Berridge, Nicola Best, Pieter Bothma, David Brealey, Robin Brittain-Long, Naomi Bulteel, Tom Burden, Andrew Burtenshaw, Vikki Caruth, David Chadwick, Duncan Chambler, Nigel Chee, Jenny Child, Srikanth Chukkambotla, Tom Clark, Paul Collini, Catherine Cosgrove, Jason Cupitt, Maria-Teresa Cutino-Moguel, Paul Dark, Chris Dawson, Samir Dervisevic, Phil Donnison, Sam Douthwaite, Ingrid DuRand, Ahilanadan Dushianthan, Tristan Dyer, Cariad Evans, Chi Eziefula, Chrisopher Fegan, Adam Finn, Duncan Fullerton, Sanjeev Garg, Sanjeev Garg, Atul Garg, Effrossyni Gkrania-Klotsas, Jo Godden, Arthur Goldsmith, Clive Graham, Elaine Hardy, Stuart Hartshorn, Daniel Harvey, Peter Havalda, Daniel B. Hawcutt, Maria Hobrok, Luke Hodgson, Anil Hormis, Michael Jacobs, Susan Jain, Paul Jennings, Agilan Kaliappan, Vidya Kasipandian, Stephen Kegg, Michael Kelsey, Jason Kendall, Caroline Kerrison, Ian Kerslake, Oliver Koch, Gouri Koduri, George Koshy, Shondipon Laha, Steven Laird, Susan Larkin, Tamas Leiner, Patrick Lillie, James Limb, Vanessa Linnett, Jeff Little, Michael MacMahon, Emily MacNaughton, Ravish Mankregod, Huw Masson, Elijah Matovu, Katherine McCullough, Ruth McEwen, Manjula Meda, Gary Mills, Jane Minton, Mariyam Mirfenderesky, Kavya Mohandas, Quen Mok, James Moon, Elinoor Moore, Patrick Morgan, Craig Morris, Katherine Mortimore, Samuel Moses, Mbiye Mpenge, Rohinton Mulla, Michael Murphy, Megan Nagel, Thapas Nagarajan, Mark Nelson, Igor Otahal, Mark Pais, Selva Panchatsharam, Hassan Paraiso, Brij Patel, Natalie Pattison, Justin Pepperell, Mark Peters, Mandeep Phull, Stefania Pintus, Jagtur Singh Pooni, Frank Post, David Price, Rachel Prout, Nikolas Rae, Henrik Reschreiter, Tim Reynolds, Neil Richardson, Mark Roberts, Devender Roberts, Alistair Rose, Guy Rousseau, Brendan Ryan, Taranprit Saluja, Aarti Shah, Prad Shanmuga, Anil Sharma, Anna Shawcross, Jeremy Sizer, Manu Shankar-Hari, Richard Smith, Catherine Snelson, Nick Spittle, Nikki Staines, Tom Stambach, Richard Stewart, Pradeep Subudhi, Tamas Szakmany, Kate Tatham, Jo Thomas, Chris Thompson, Robert Thompson, Ascanio Tridente, Darell Tupper-Carey, Mary Twagira, Andrew Ustianowski, Nick Vallotton, Lisa Vincent-Smith, Shico Visuvanathan, Alan Vuylsteke, Sam Waddy, Rachel Wake, Andrew Walden, Ingeborg Welters, Tony Whitehouse, Paul Whittaker, Ashley Whittington, Meme Wijesinghe, Martin Williams, Lawrence Wilson, Sarah Wilson, Stephen Winchester, Martin Wiselka, Adam Wolverson, Daniel G. Wooton, Andrew Workman, Bryan Yates, and Peter Young.

---

**Supplementary Figure 1.**
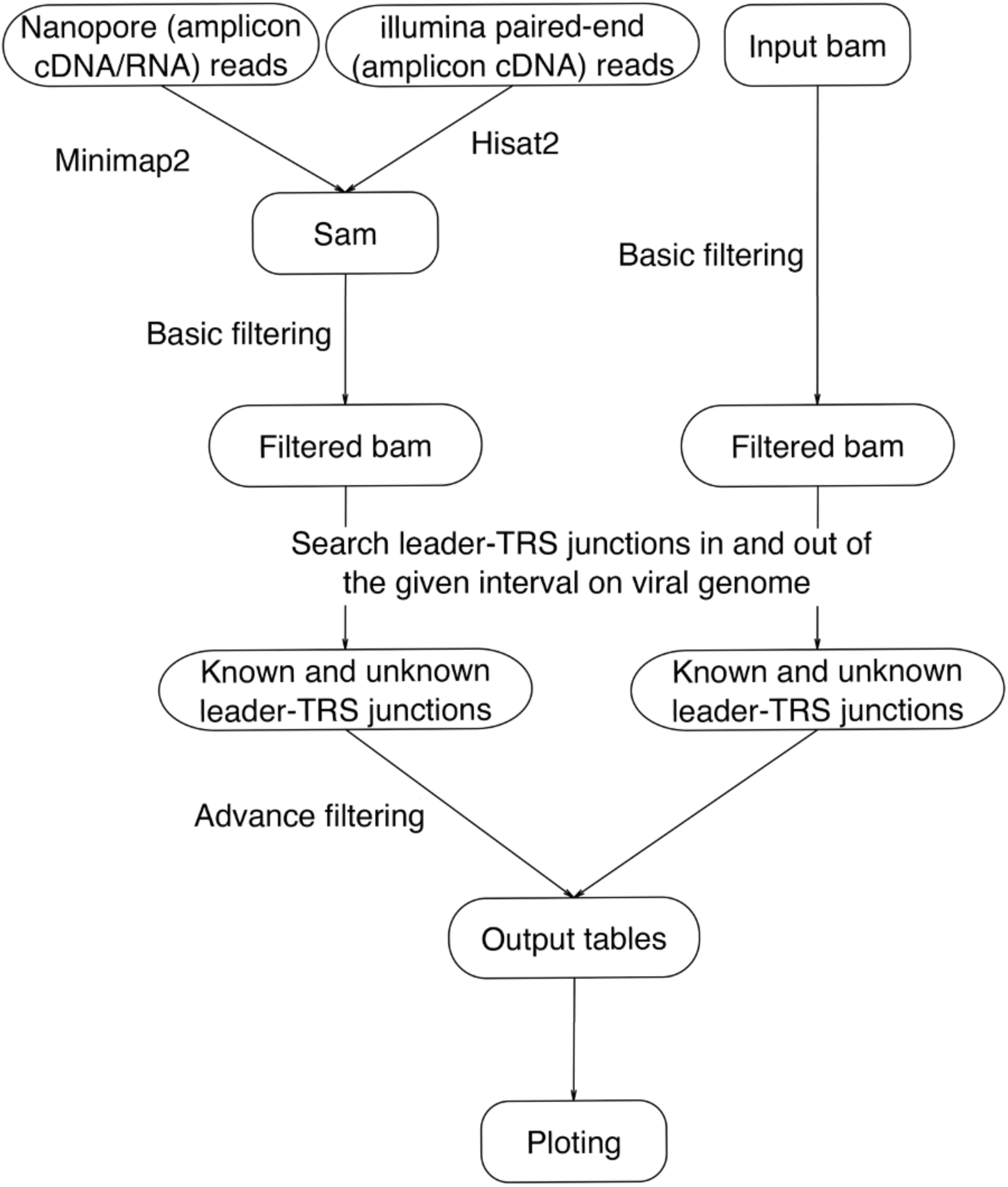
Bioinformatics pipeline for the identification of leader-TRS junctions in sequencing data from SARS-CoV-2 infected material with LeTRS. This can be rapidly adapted for other coronaviruses. LeTRS can work from Nanopore or Illumina amplicon data or more unbiased approaches such as metagenomic or Illumina sequencing by using a BAM file.

**Supplementary Figure 2.**
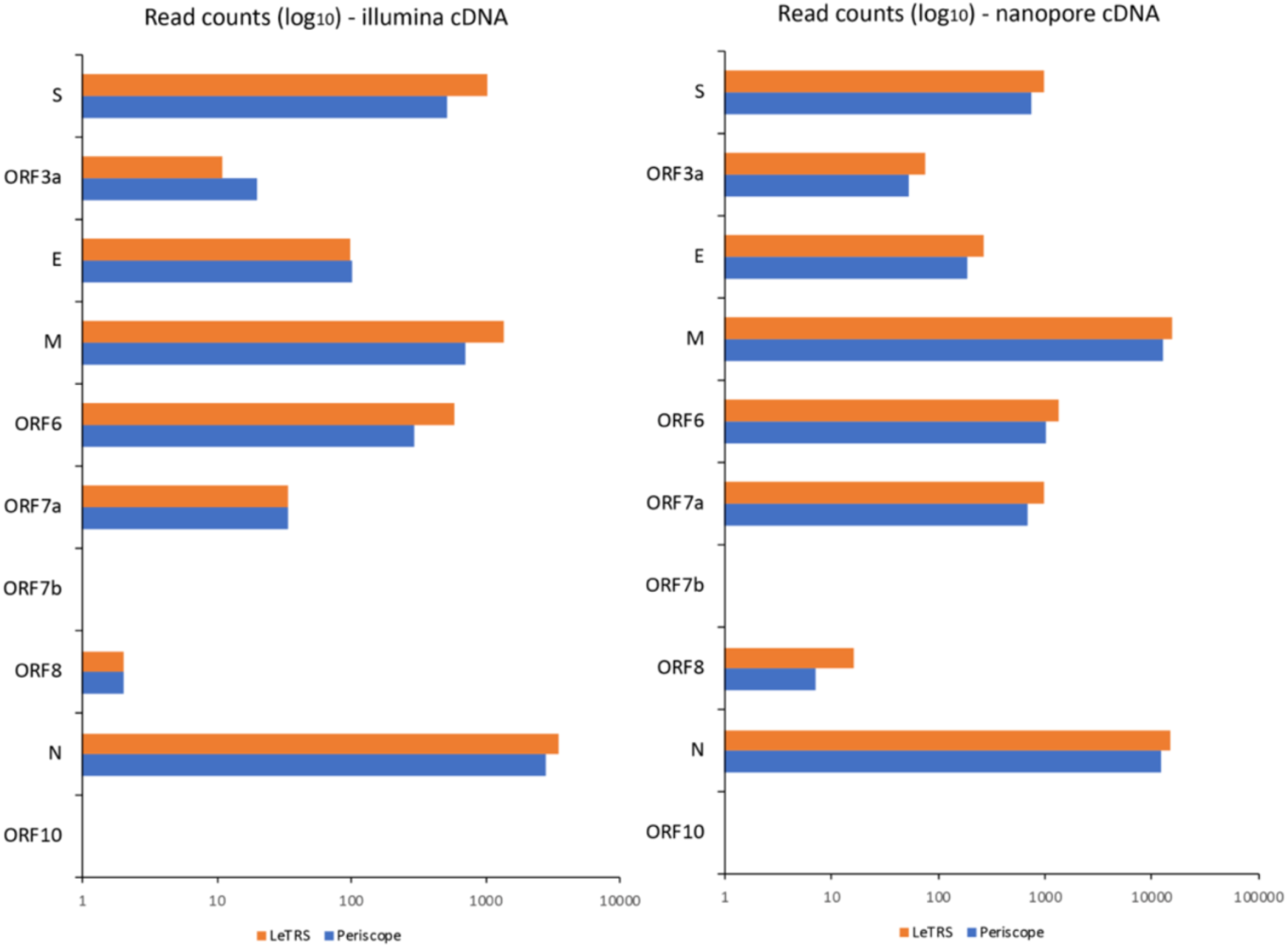
Comparison of LeTRS to Periscope with the Illumina (left) and Nanopore (right) ARTIC amplicon sequencing test data sets by using the number of reads with at least one primer sequences at either end in LeTRS and the number of “High Quality” reads (the reads with both 32 nts leader sequences and known TRS-orf boundary) in Periscope.

**Supplementary Figure 3.**
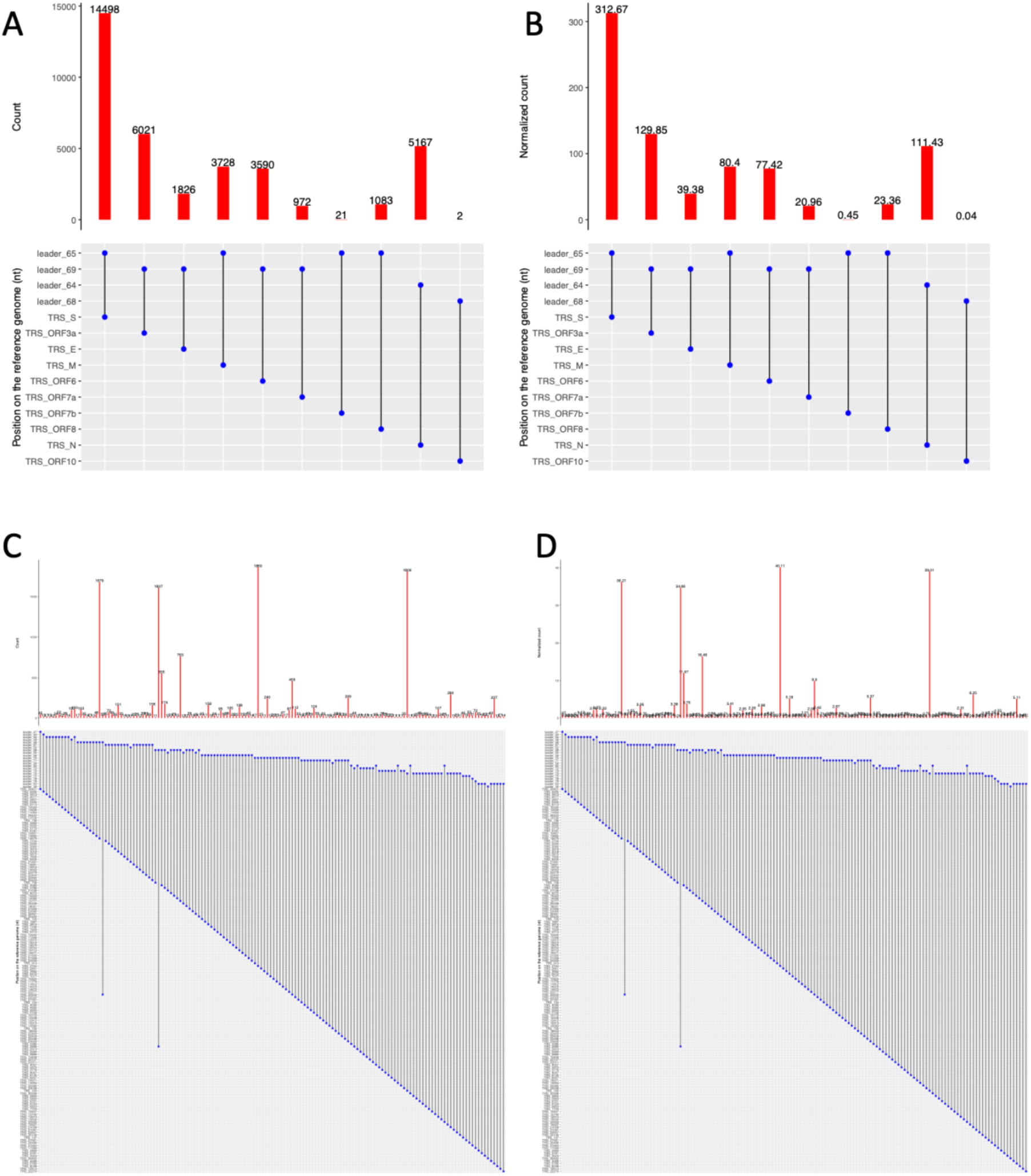
Raw (A and C) and normalised (B and D) expected (upper) and novel (lower) leader-TRS gene junctions count in the infecting SARS-CoV-2 inoculum source used for NHP study, sequenced by Illumina ARTIC method (Supplementary Table 8).

**Supplementary Figure 4.**
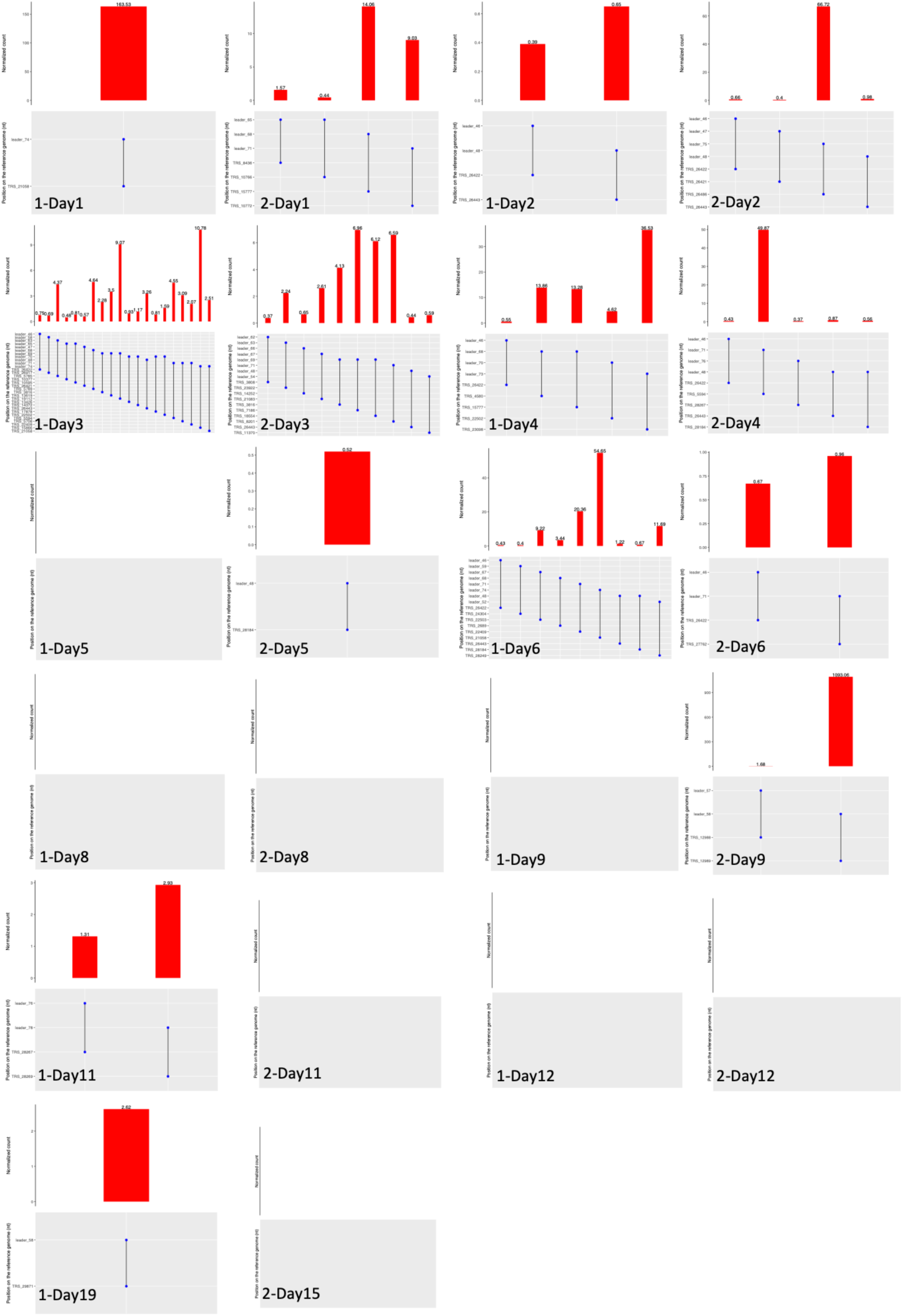
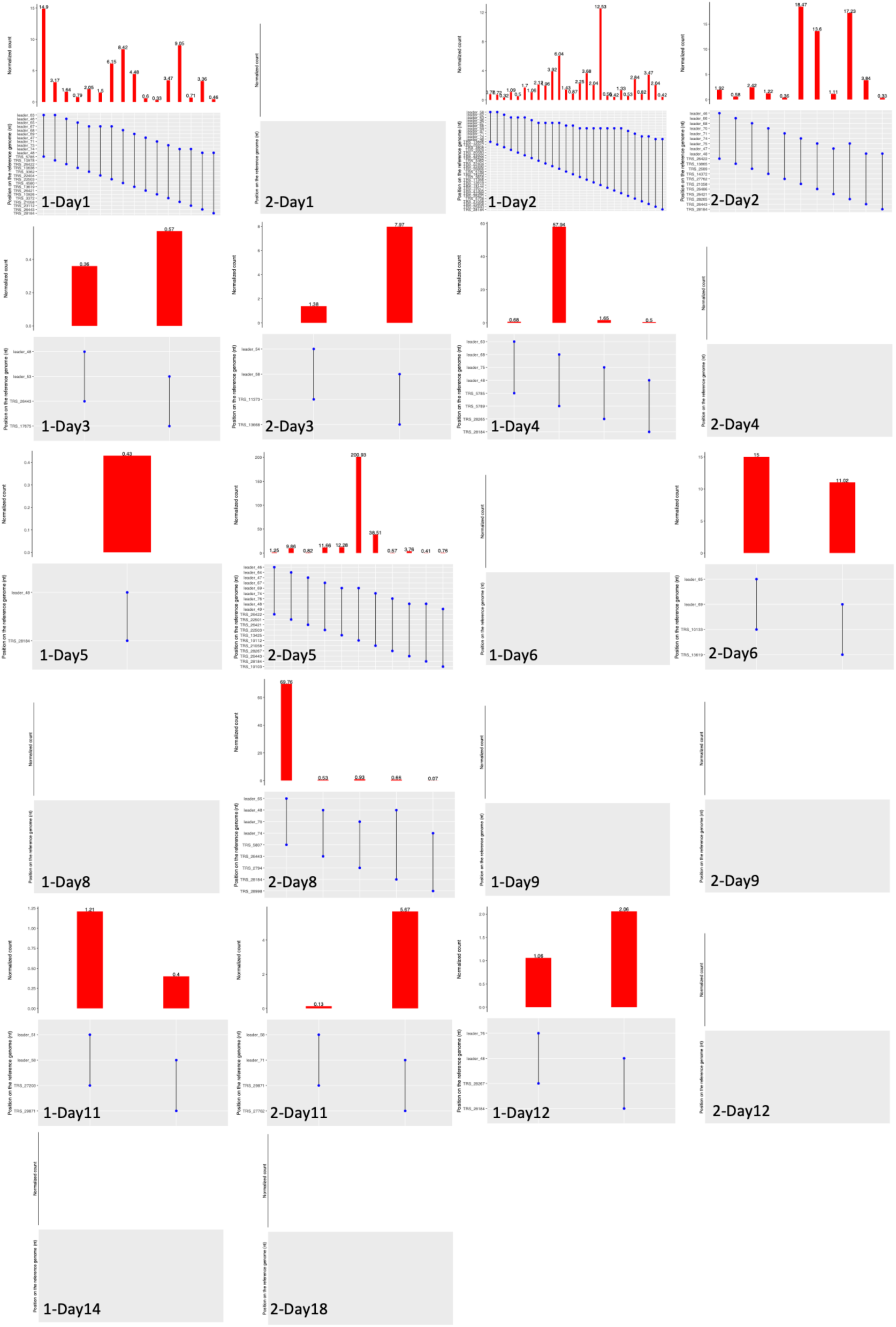
Novel leader-TRS gene junctions identified for cynomolgus macaques (Supplementary Table 9). The number before “-Day” indicated the group of cynomolgus macaques.

**Supplementary Figure 5.**
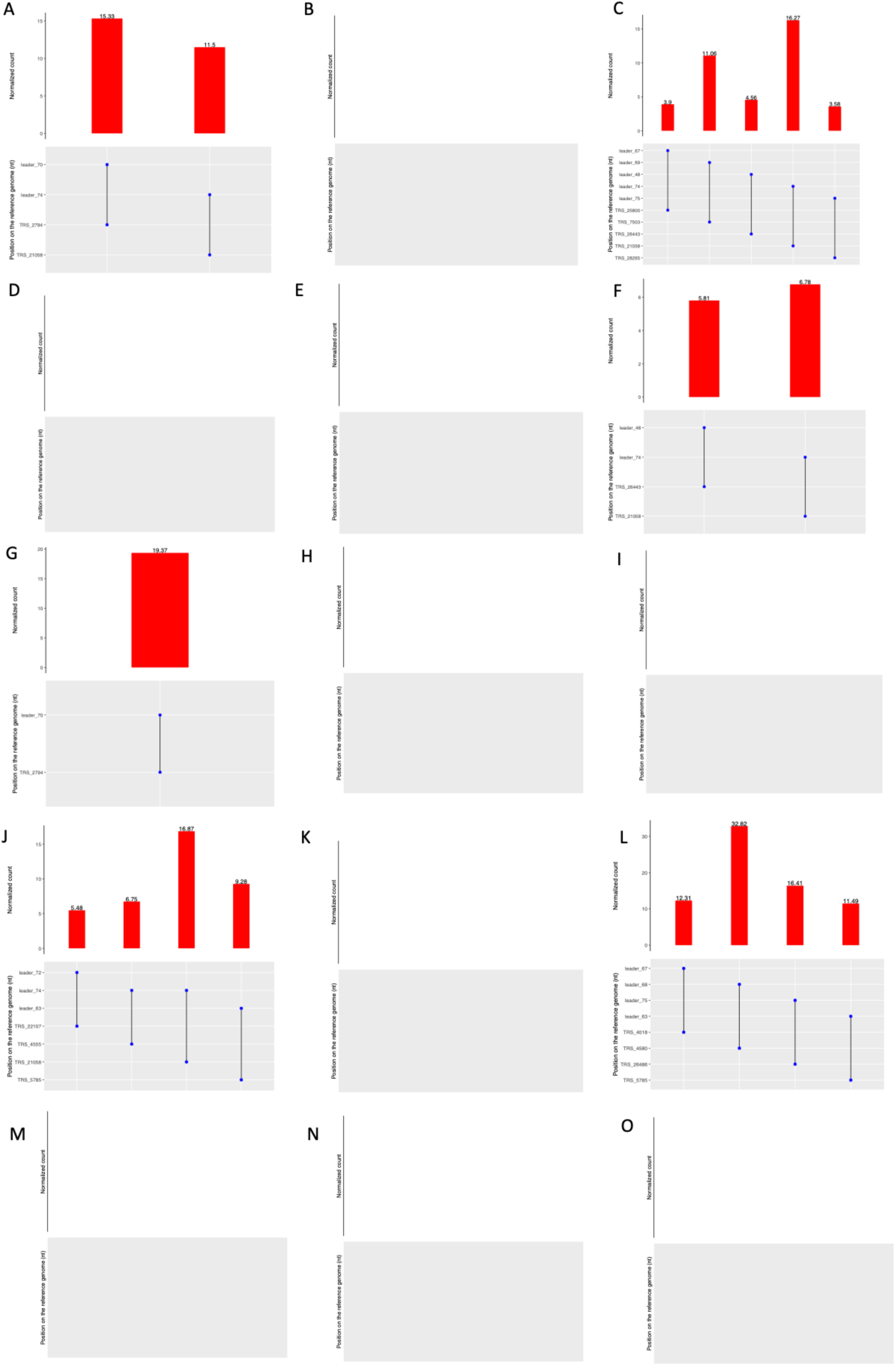
Novel leader-TRS gene junctions identified in nasopharyngeal swabs from human patients sequenced using the ARTIC-Illumina approach (Supplementary Table 7).

**Supplementary Figure 6.**
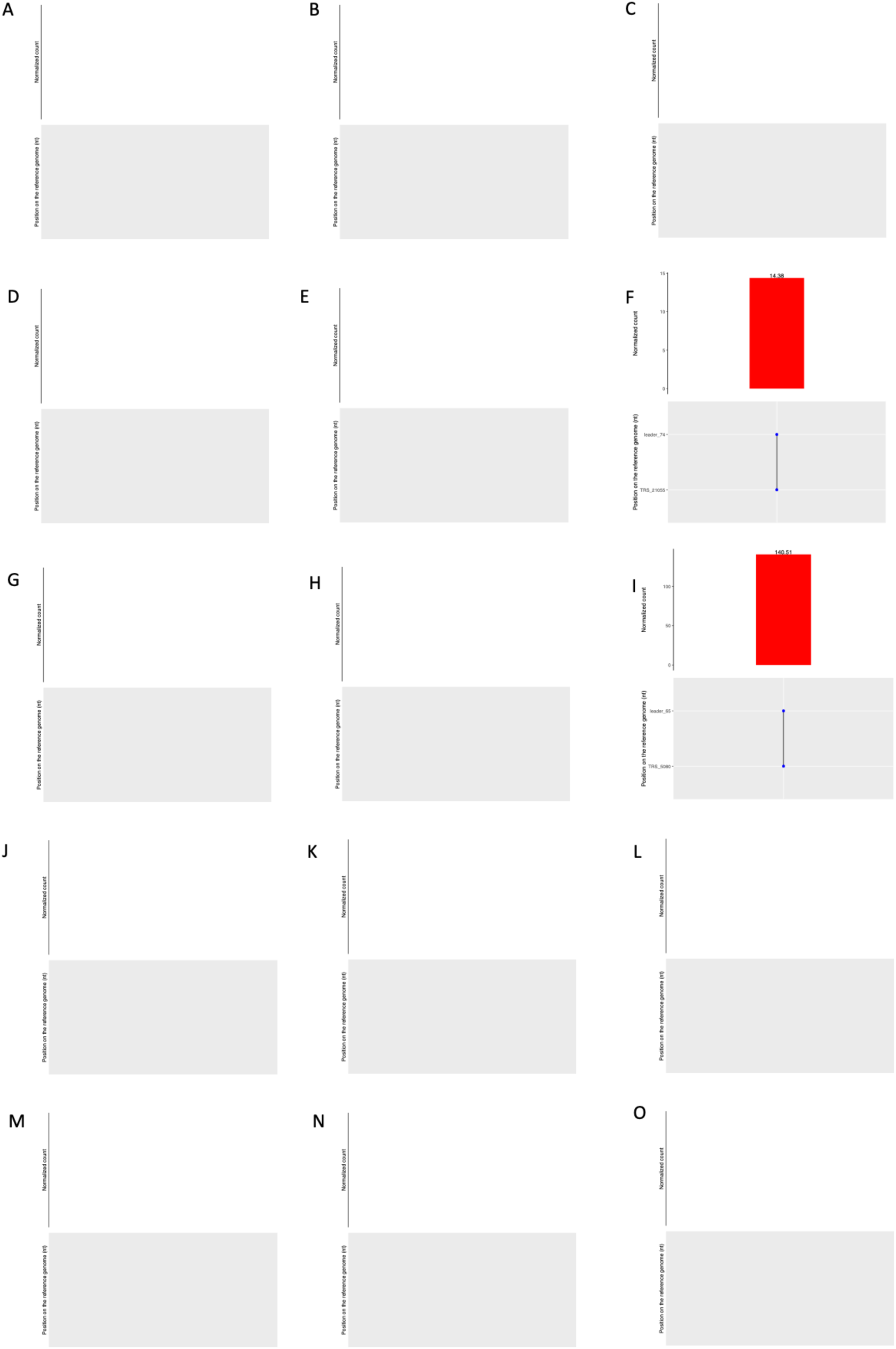
Novel leader-TRS gene junctions identified in nasopharyngeal swabs from human patients sequenced using the ARTIC-Nanopore approach (Supplementary Table 10).

## Supplementary data

Table S1. The LeTRS output table for details of known sgmRNA in the tested Nanopore ARTIC v3 primers amplicon sequencing data. “peak_leader” and “peak_TRS_start” point to the leader-TRS junctions in Table 1, “ACGAAC” indicates if there is a ACGAAC sequence in the “TRS_seq” (TRS sequences), “20_leader_seq” refers to the 20 nucleotides before the end of leader, and “AUG_postion” and “first_orf_aa” refer to the first AUG position and translated orf of the sgmRNA.

Table S2. The LeTRS output table for details of novel sgmRNA in the tested Nanopore ARTIC v3 primers amplicon sequencing data. “peak_leader” and “peak_TRS_start” point to the leader-TRS junctions in Table 2, “ACGAAC” indicates if there is a ACGAAC sequences in the “TRS_seq” (TRS sequences), “20_leader_seq” refers to the 20 sequences before the end of the leader, “AUG_postion” and “first_orf_aa” refer to the first AUG position and translated orf of the sgmRNA, and “known_ATG” indicates if the first AUG position is the same as a known sgmRNA.

Table S3. The LeTRS output table for details of known sgmRNA in the tested Nanopore direct RNA sequencing data. “peak_leader” and “peak_TRS_start” point to the leader-TRS junctions in Table 3, “ACGAAC” indicates if there is a ACGAAC sequence in the “TRS_seq” (TRS sequences), “20_leader_seq” refers to the 20 nucleotides before the end of leader, and “AUG_postion” and “first_orf_aa” refer to the first AUG position and translated orf of the sgmRNA.

Table S4. The LeTRS output table for details of novel sgmRNA in the tested nanopore direct RNA sequencing data. “peak_leader” and “peak_TRS_start” point to the leader-TRS junctions in Table 4, “ACGAAC” indicates if there is a ACGAAC sequences in the “TRS_seq” (TRS sequences), “20_leader_seq” refers to the 20 nucleotides before the end of leader, “AUG_postion” and “first_orf_aa” refer to the first AUG position and translated orf of the sgmRNA, and “known_AUG” indicates if the first AUG position is the same as a known sgmRNA.

Table S5. The LeTRS output table for details of known sgmRNA in the tested Illumina ARTIC v3 primers amplicon sequencing data. “peak_leader” and “peak_TRS_start” point to the leader-TRS junctions in Table 7, “ACGAAC” indicates if there is a ACGAAC sequence in the “TRS_seq” (TRS sequences), “20_leader_seq” refers to the 20 nucleotides before the end of leader, and “ATG_postion” and “first_orf_aa” refer to the first AUG position and translated orf of the sgmRNA.

Table S6. The LeTRS output table for details of novel sgmRNA in the tested Illumina ARTIC v3 primers amplicon sequencing data. “peak_leader” and “peak_TRS_start” point to the leader-TRS junctions in Table 8, “ACGAAC” indicates if there is a ACGAAC sequences in the “TRS_seq” (TRS sequences), “20_leader_seq” refers to the 20 nucleotides before the end of leader, “AUG_postion” and “first_orf_aa” refer to the first AUG position and translated orf of the sgmRNA, and “known_AUG” indicates if the first AUG position same as a known sgmRNA.

Table S7. Leader-TRS gene junctions of reads with at least one primer sequence derived from sequence data from 15 human patients sequenced with the ARTIC pipeline via Illumina.

Table S8. Leader-TRS gene junction count in the infecting SARS-CoV-2 inoculum source used for the NHP study, sequenced by Illumina ARTIC method.

Table S9. Analysis of leader TRS-gene junction, abundance of reads with at least one primer sequence at either end in longitudinal nasopharyngeal samples taken from two non-human primate models (cynomolgus and rhesus macaques) of SARS-CoV-2 in groups. SARS-CoV-2 was amplified using the ARTIC approach and sequenced by Illumina. The data is organised into groups of animals for the cynomolgus macaque groups 1 and 2 that were with “-1” and “-2” in the excel sheets.

Table S10. leader-TRS gene junctions of reads with at least one primer sequence derived from sequence data from 15 human patients sequenced with the ARTIC pipeline via Nanopore.

Table S11. Novel leader-TRS junctions centred around the known gene open reading frame but out of the search interval in the analysis of cell culture, non-human primate and human sequencing data.

